# Drift-diffusion dynamics of hippocampal replay

**DOI:** 10.1101/2025.10.14.682470

**Authors:** Zhongxuan Wu, Xue-Xin Wei

## Abstract

Replay in the hippocampus during sharp-wave ripples is thought to play major roles in learning and memory. However, existing analysis methods often lead to inconsistent and inaccurate metrics for characterizing replay dynamics. We develop a novel computational framework that model the replay dynamics using drift-diffusion process. Further, to capture the potentially rich population-level dynamics during sharp-wave ripples, our model allows switching between multiple well-motivated types of dynamics. Applications of our method to rat hippocampal recordings lead to insights into a number of important open questions, including: (i) whether the speed of most replay events is comparable to real-world running; (ii) whether replays follow a random walk; (iii) whether “preplay” events exist. Overall, our approach enables precise characterizations and unambiguous interpretations of population dynamics during sharp-wave ripples, which more broadly can provide a better understanding of the functions and underlying mechanisms of replay.

## 1 Introduction

Neural activities in the mammalian hippocampus during sharp-wave ripples (SWRs) have been found to “replay” previous experience in forward [1, 2] or reverse [3, 4] order, both during sleep [5–8] and awake immobility [3, 9]. Replays in the hippocampus are also found to be coordinated with neural activities in other brain regions, e.g., prefrontal cortex [10–12], visual cortex [13], and brain-wide spontaneous dynamics [14]. While initially reported in rodents, experimental evidence for replay in the human brain has also been reported in recent studies [15–19]. As a form of selective and experience-dependent neural activity, replays have been proposed to play an important role in learning, memory consolidation, and planning [20–23], although their precise functions remain to be determined.

Surprisingly, even given decades of research on hippocampal replay, the methodology of detecting replays and, at the same time, identifying their properties on an event-by-event basis remain an ongoing challenge, both statistically and also with respect to defining the phenomenon [20, 24]. Most studies rely on template matching, where putative replay activity is compared to a template generated from the population activity during “online” periods, typically taken to be running [5, 6]. More specifically, this template can be defined using either a sequential order of neurons or the sequential structure in a decoded variable, conventionally spatial location. However, many SWR events remain ambiguously classified according to this approach, rendering a challenge for subsequent analyses and, further, suggesting that many replays may have structure that is not well-captured by existing approaches. As an example, in prior methods, the speed and quality (coherence) of replays are entangled. Replay events with high speed tend to be significant despite having lower coherence, even though these variables are ideally independently estimated. Indeed, recent studies suggest that replays may exhibit richer dynamics than traditionally considered or assumed [25, 26], and the properties of replay dynamics may have major implications for learning [27]. In this way, the lack of a principled and appropriately general computational framework for characterizing the dynamics of replay constitutes a major obstacle to analyzing and interpreting ongoing experimental data and findings, respectively.

Here we address this gap by developing a principled probabilistic approach for unbiased characterization of the dynamics of replay. This method builds upon prior work [25, 26, 28–30] and overcomes multiple limitations in accurate identification of replay dynamics. We will show that applications of our modeling framework to neural population recordings from the rodent hippocampus lead to new insights into the precise neural dynamics of the hippocampal replay. First, in contrast to recent observations in [25], our results show that only a small fraction of SWRs (∼ 5%) are stationary, and furthermore, those with drift-diffusion dynamics generally have a much higher speed than the animal’s movement. Second, we find that replay dynamics in 1-D environments are heterogeneous, and are inconsistent with a random walk model proposed previously [31]. The mean squared displacement of replay trajectories scales quadratically with time, supporting the presence of substantial drifts. Third, we find that only a tiny fraction of SWR events (less than 1%) exhibit sequential structure at the timescale of 100 ms before the animal had spatial experience in an environment. This suggests that, while neural activity in the hippocampus may be coordinated before the animal’s spatial experience (i.e., preplay [32]), the level of coordination is much weaker than that after spatial experience. By enabling more flexible and interpretable characterization of replays, this new analysis framework may provide a path toward better understanding of the functions and mechanisms of replay.

## 2 Results

A replay event is generally thought to be a sequence of neural activity that encodes a traversal of a particular state space (e.g., positions on the track). Most previous studies use a two-step procedure to analyze replays: (i) decode spatial location from neural population activity, typically using a Bayesian decoder that yields decoded posterior probabilities; (ii) apply a “manually” specified form of template matching to the decoded posterior probabilities to detect putative sequential structure that would qualify as a (traditional) replay. Despite the success of this approach in identifying classical replays, the major limitation is the rigidity of the pre-specified template, which effectively disregards the potential richness of replay dynamics. A second major limitation is that the estimation in the second step fails to incorporate uncertainty, which limits quantitative evaluation of robustness. To address these issues, we develop a novel set of state-space models based on drift-diffusion dynamics to characterize the structure of replay events. By basing dynamics on drift-diffusion, a general yet interpretable and parsimonious process, our approach rigorously formalizes past intuition regarding replays, and moreover allows for direct probabilistic inference of replay dynamics from neural data.

### 2.1 Drift-diffusion models of replay dynamics

Consider an experiment where the animal is running on a one-dimensional track (Fig. 1A,B), a common behavioral paradigm in studying replays. Although we focus on modeling replays in 1-D environment, our model can be generalized to more complex environments. We assume that hippocampal activity during a replay event encodes a sequence of positions on the track (Fig. 1C). However, in contrast to previous work, we propose to model changes in encoded positions during replays as evolving according to the dynamics specified by drift-diffusion. Formally, these dynamics of replayed positions *z* during putative replays (population activity during SWRs) evolves according to (Fig. 1D):

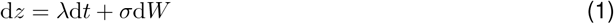

where *λ* and *σ* are the drift and diffusion parameters, respectively. d*W* denotes the standard Brownian increment. This model can be implemented in discrete time steps, with the dynamics described by the state-transition matrix. We further assume that, given a latent state, the activity of individual neurons follows an independent Poisson process with firing rate dependent on location-specific fields (place fields) –an assumption shared with the conventional Bayesian decoding method for analyzing replays (see Methods).

**Figure 1:**
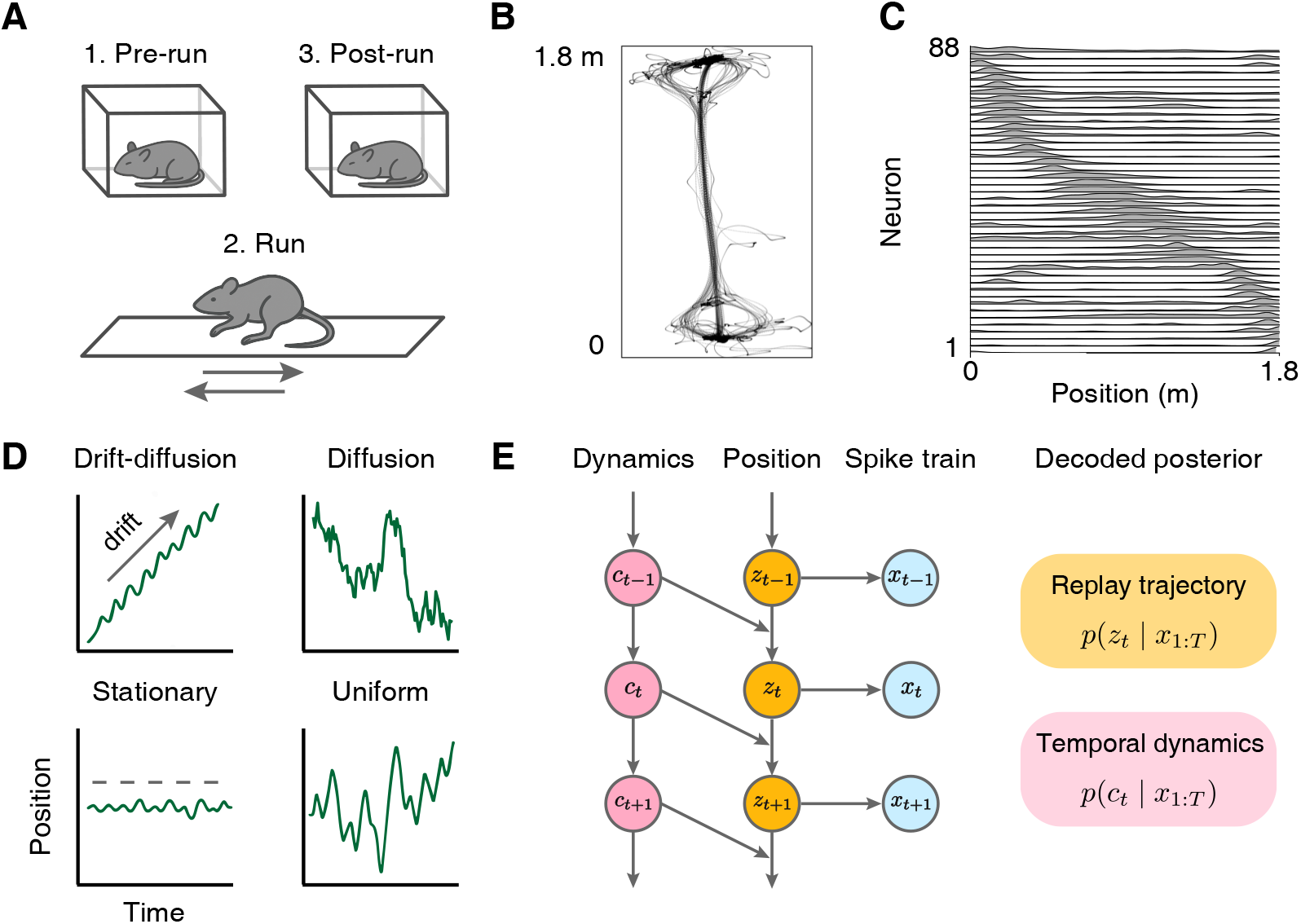
Animal behavior and the overall computational modeling framework. **A.** Schematic diagram of a commonly used behavioral paradigm for studying hippocampal replay. Each recording session consists of three behavioral epochs: a pre-run sleep epoch in an isolated box, an active run epoch on a 1-D linear track, and a post-run sleep epoch in the same box used for the pre-run. **B**. Head-location tracking data during the run epoch from an example recording session. The animal performed repeated back-and-forth traversals along the 1.8 m track. The track is 6.35 cm wide, with two ends expanding to 12.7 cm for reward wells. Data replotted from [33]. **C**. Place fields of the recorded CA1 neurons, estimated from spikes during locomotion (> 5 cm/s) using 2 cm spatial bins and Gaussian smoothing (*σ* = 5 cm). Only half of the recorded population is plotted for clarity. Data from [33]. **D**. Potential spatiotemporal dynamics underlying the latent trajectory during sharp-wave ripples (SWRs). *Drift-diffusion*: a mixture of deterministic drift and stochastic diffusion, *dz* = *λ dt* + *σ dW*, where the drift parameter *λ* sets the “speed” and diffusion parameter *σ* sets the amplitude of Brownian motion fluctuations. *Diffusion*: pure Brownian motion, *dz* = *σ dW*, representing an unbiased random walk. *Stationary* : *dz* = 0, indicating no movement over time. *Uniform*: drift-diffusion process with near-zero *λ* and large *σ*, implemented as independent resampling of position from a uniform distribution at each time step. Note that the uniform trajectory has no temporal continuity. **E**. Left, probabilistic graphical model of a switching HMM, where a discrete variable of dynamics (regime) *c*_*t*_ determines a regime-specific transition probability for latent position *z*_*t*_. *z*_*t*_ then generates the observed spike counts *x*_*t*_ with Poisson emissions parameterized by estimated place fields. Right, applying probabilistic inference to the switching HMM produces a marginal posterior of trajectory *p*(*z*_*t*_ | *x*_1:*T*_) in conjunction with a marginal posterior of regime variable *p*(*c*_*t*_ | *x*_1:*T*_). For derivations of marginal posteriors, see Methods and Supplementary Information.

In this framework, the expected latent state at the next time step is determined by the current state and the drift rate, with the variability of the states determined by the diffusion parameter. In this way, both parameters (drift and diffusion) are highly interpretable. The drift parameter is analogous to the speed (or “virtual speed”) of the replay event. The diffusion parameter quantifies to how noisy the replay trajectory is. Different values of two parameters can capture a variety of dynamical patterns while retaining interpretability (Fig. 1D). In the case of zero drift and nonzero diffusion, the latent trajectory is equivalent to one-dimensional Brownian motion, a pattern recently proposed to capture replay dynamics [31]. In the case where both drift and diffusion parameters are near zero, the trajectory is largely stationary, a dynamical pattern emphasized in [25]. Finally, in the case where the drift rate nearly zero, and diffusion is large, the discretized trajectory would behave similarly to a collection of independently sampled states having little coherence. We refer to this type of event as a uniform event. Conceptually, this is the opposite regime of replay events discussed in previous studies.

#### Extended modeling framework to incorporate switching dynamics

A fundamental matter that remains unresolved regarding replays is whether and to what extent individual SWR events express multiple dynamical patterns serially [25]. To capture the potential richness of the dynamics of an SWR event, we incorporate the switch between different types of dynamics within an individual event (Fig. 1E). Importantly, both the type of dynamics and the states (i.e., positions) can be inferred from the neural activity based on our model. The resulting one-step framework represents the most powerful way to date to clarify whether and to what extent there are mixtures of dynamics. Specifically, we propose that a replay event may be serially comprised of three types of dynamics (regimes): drift-diffusion, stationary, and uniform sampling (Fig. 2A). For a given event, in addition to the latent position evolution, there is also a finite probability of switching to a different type of dynamics. We assume that the transition probability matrix between regimes has equal self-transition probability *q* and identical switching probabilities 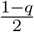 of switching to the other two regimes. The parameter *q* is optimized for individual replay events and thus can capture variability of dynamical switching which may be expressed across different events, as suggested in previous work [35]. The transition matrix between different regimes can be easily incorporated into the model by conditioning the state transition matrix for each type of dynamics. We develop inference procedures for estimating the parameters from the neural population activity, leveraging established techniques in HMMs and state-space models [36–43]. For each SWR event, we optimize the model parameters {*λ, σ, q*} by maximizing the likelihood of observing the neural population activity. With optimized parameters, we can calculate the posterior distribution of dynamics and positions (see Methods and Supplementary Information A for details). Fig. 2B-H show example events from analyzing a hippocampal population recording dataset [33].

**Figure 2:**
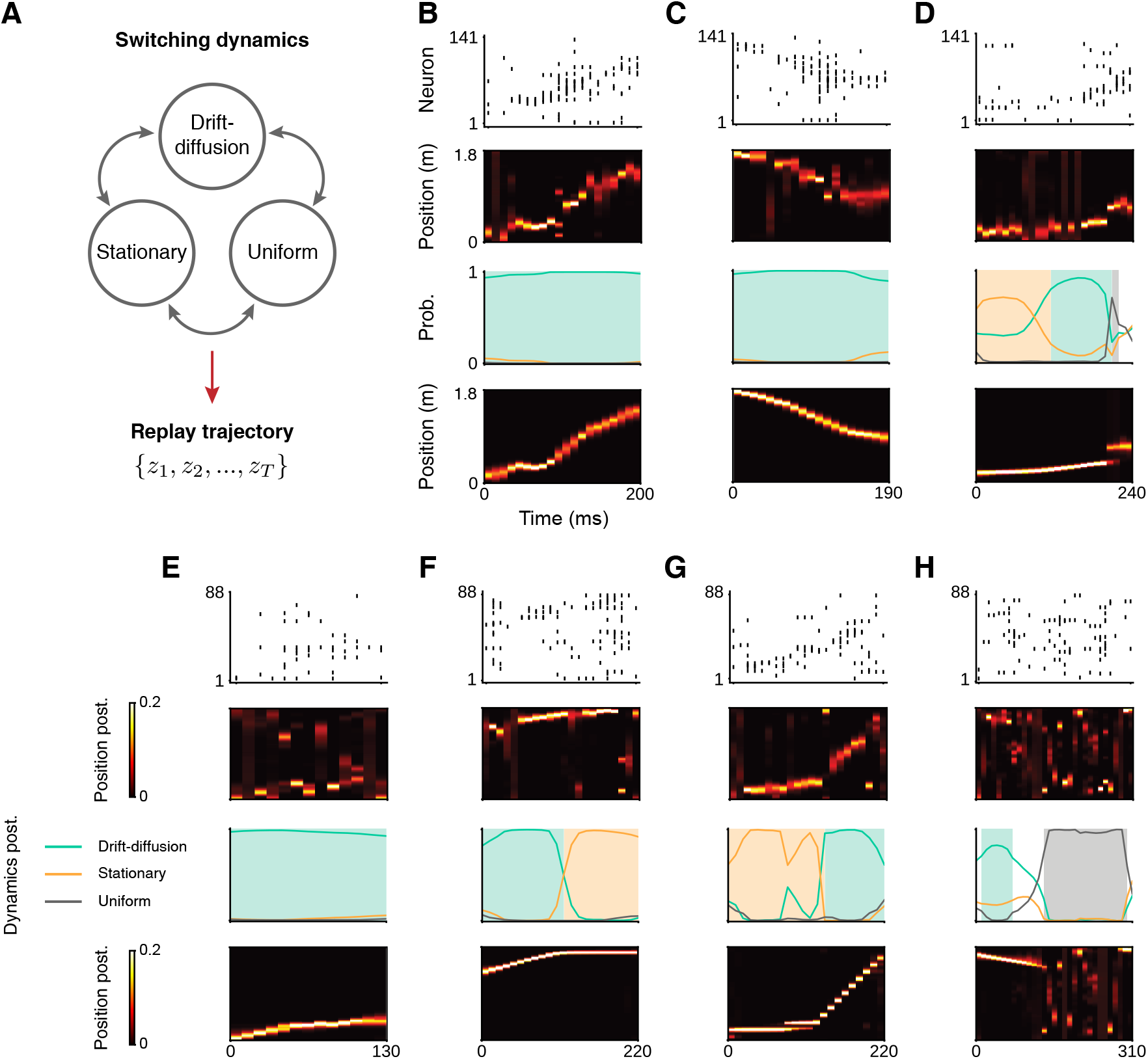
Decoding the heterogeneous dynamics in rat hippocampal SWRs using switching HMMs. **A.** Schematic diagram of the switching dynamics in our framework. At each time step, the model can switch between three basic dynamics—drift-diffusion, stationary and uniform (see definitions in Fig. 1D). The probabilities of these basic dynamics together determine the temporal transition of latent positions *p*(*z*_2_ | *z*_1_), *p*(*z*_3_ | *z*_2_), …, *p*(*z*_*T*_ | *z*_*T* −1_). **B-H**. Seven representative SWRs from a rat hippocampal recording dataset [33] are each shown as four subpanels. Top to bottom: (1) Spike raster of the SWR. Spikes are binned using 10 ms non-overlapping time bins. For visualization, place cells are sorted by the peak location of their place fields. (2) Posterior probability of the latent positions decoded from a memoryless Bayesian decoder [34]. (3) Posterior probability of the three basic dynamics decoded from our model, *p*(*c*_*t*_ | *x*_1:*T*_). Background shading indicates the assigned type of dynamics based on the posterior. If *p*(*c*_*t*_ = any type of dynamics | *x*_1:*T*_) *>* 0.7, we assign that type at time step *t*. Otherwise, if *p*(*c*_*t*_ = drift-diffusion) + *p*(*c*_*t*_ = stationary) *>* 0.7, we label such time steps stationary, as drift-diffusion reduces to stationary in this case. Time steps that do not meet any of these criteria are labeled unclassified. Color code: drift-diffusion, green; stationary, orange; uniform, gray; unclassified, none. (4) Posterior probability of the latent positions decoded from our model, *p*(*z*_*t*_ | *x*_1:*T*_).

### 2.2 Validation of the modeling framework

Our full model of replay dynamics can be interpreted as a specific type of switching HMM. Although identifiability has been established for related models [44, 45], two potential issues are relevant for our formulation: (i) under maximum-likelihood estimation (MLE), are the drift and diffusion parameters governing the latent-position transition probability recoverable, and (ii) can dynamic regimes be correctly classified? A practical constraint is that replay events are brief with a typical length of 100-250 ms and recordings comprise only dozens to a few hundred place cells. This relatively limited quantity of data may make estimation challenging. Thus, it is materially important to evaluate the performance of our inference procedures under conditions that are relevant to this experimentally relevant constraint.

We first validated parameter identifiability for the basic drift-diffusion model without switching dynamics.

We generated latent position trajectories from a two-dimensional grid of drift and diffusion values, then simulated spike trains from synthetic place fields conditioned on each trajectory. By fitting our model to these synthetic data, we can assess how well the model parameters can be recovered (Fig. 3A). In general, both drift and diffusion parameters were satisfactorily recovered (Fig. 3B-C). We found that the diffusion parameter was considerably more variable than the drift, and variance of the diffusion parameter tended to increase with the true diffusion magnitude. We also found that the diffusion estimates tend to be underestimated because each replay event only contains a limited number of time steps (see SI for a further discussion on the origin of this estimation bias).

**Figure 3:**
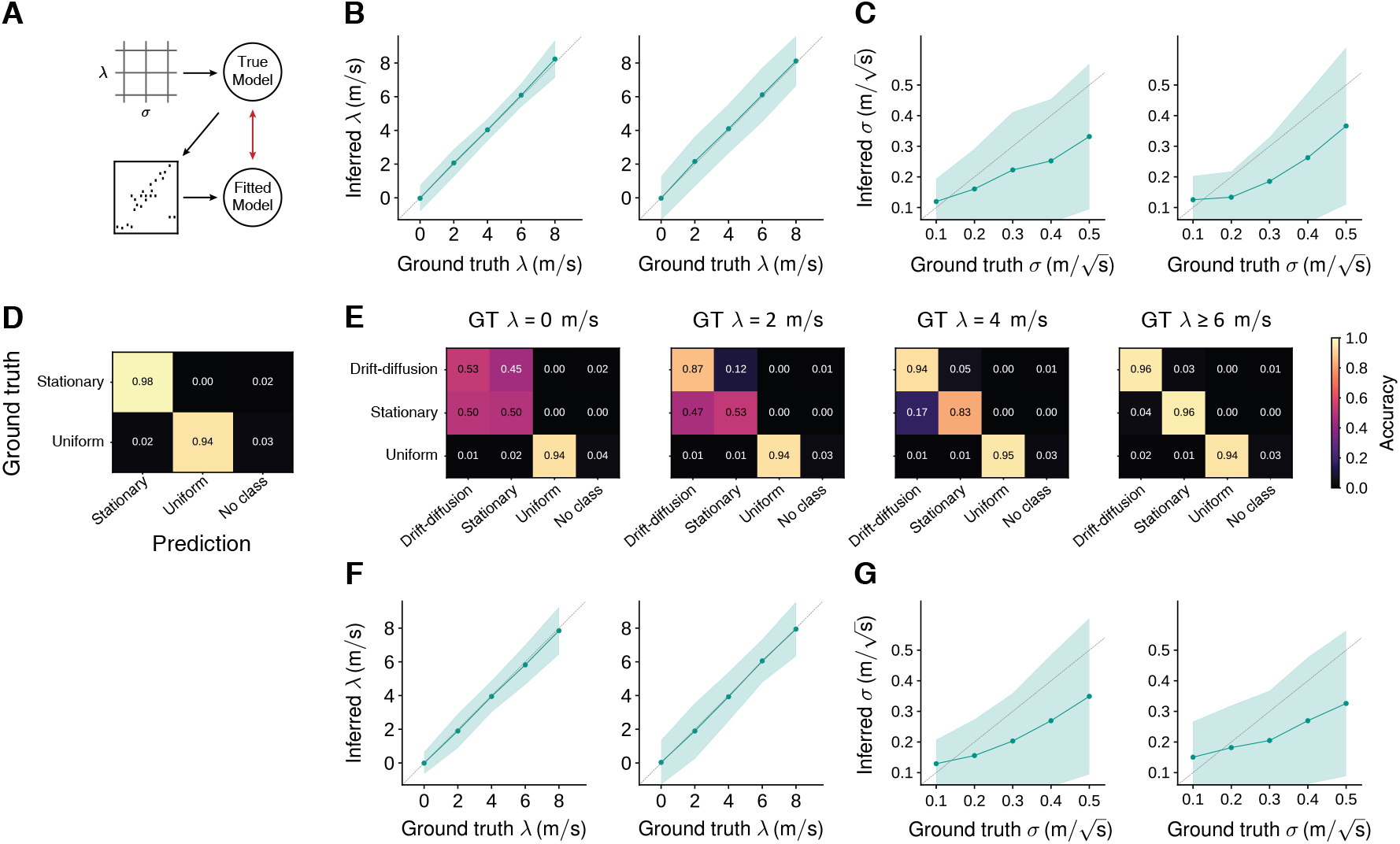
Identifiability of the model under realistic experimental conditions. **A.** Schematic diagram of the simulation pipeline. We considered two simulation settings: pure drift-diffusion (**B-C**), in which only drift-diffusion regime was included, and regime switching (**D-G**), in which the ground-truth dynamics could switch between drift-diffusion, stationary, and uniform regimes. In the pure drift-diffusion setting, ground-truth drift-diffusion parameter pairs (*λ, σ*) were first sampled on a grid. The sampled parameters determine the drift-diffusion dynamics which was used to generate multiple trajectories on a 2 m linear track. Each trajectory was then used to generate one 150 ms SWR event with population activity of *N* = 120 simulated place cells whose place fields uniformly tile the track. In the regime-switching setting, each SWR consisted of two concatenated 150 ms parts drawn from different regimes. Within each part, trajectories were generated according to the sampled regime. If the regime is drift-diffusion, the data generation followed the same procedure as in the pure drift-diffusion setting. If the regime is stationary or uniform, trajectories were generated using their corresponding dynamics and population activities were generated using the same place cell population. Each simulated SWR event was analyzed with our inference pipeline to obtain marginal posterior of dynamics and, if the ground-truth regime is drift-diffusion, parameter estimates 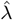 and 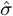. The marginal posterior of dynamics was classified into a discrete regime sequence {*ĉ*_*t*_} following the convention in Fig. 2B-H. The inferred 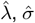 and {*ĉ*_*t*_} were compared with the ground truth to quantify recovery performance. **B**. The parameter recovery of *λ* when *σ* is held at 0.2 (left) or 0.4 (right) 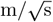. **C**. The parameter recovery of *σ* when *λ* is held at 0 (left) or 2 (right) m/s. Solid lines show the mean over 100 repeats and shaded bands span *±*1 standard deviation. The diagonal line denotes perfect recovery. **D**. Confusion matrix of regime classification accuracy for simulated SWRs when drift-diffusion regime is not involved. **E**. Confusion matrices of regime classification accuracy for simulated SWRs when drift-diffusion regimes with different ground-truth (GT) drift values are involved. From left to right, the ground truth drift is 0, 2, 4, ≥ 6 (m/s), respectively. Color intensity reflects the classification accuracy. **F**. The parameter recovery of *λ* when *σ* is held at 0.2 (left) or 0.4 (right) 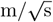. **G**. The parameter recovery of *σ* when *λ* is held at 0 (left) or 2 (right) m/s. Solid lines show the mean over 100 repeats and shaded bands span *±*1 standard deviation. The diagonal line denotes perfect recovery.

We next evaluated identifiability in the switching HMM, the model that enables inference of switching between dynamical regimes. As before, we simulated latent trajectories with regime switches and generated spike trains from synthetic place fields, then optimized the full model. We first asked whether dynamics classes could be correctly decoded. We found that fitting yielded robust regime classification overall, as shown by confusion matrices comparing inferred and true regimes for simulated SWRs (Fig. 3D-E). As expected, when both drift and diffusion are near zero and observed replay sequence is short, the separability between drift-diffusion and stationary regimes is reduced. With a substantial drift magnitude, separability improves and classification accuracy between drift-diffusion and stationary increases markedly (Fig. 3E). Consistent with the basic drift-diffusion analysis, the full model also yields accurate parameter recovery. MLE estimates of both drift and diffusion recover the ground truth across simulated conditions reasonably well (Fig. 3F-G). However, there did remain modest underestimation of the diffusion parameter, attributable to short events. Together, these results show that our full model can reliably identify the dynamical regimes and the drift-diffusion parameters under experimentally realistic data sizes.

#### Comparison to prior methods

We compared our modeling framework with the traditional 2-step analysis procedure, which first uses Bayesian decoding to map the neural population activity onto the posterior distribution over positions [34], then estimates descriptive statistical parameters (such as absolute correlation, slope of replay trajectory, maximum jump distance) [46, 47] from the resulting posteriors (see Fig. S1A). Using simulated data, we compared the drift/diffusion parameters to the key metrics used in prior studies. Based on extensive simulations, we found that, while the three key metrics may capture some structure in replays as conceived when they were developed, each measure is relatively confounded with the estimated speed and quality of the candidate replay event (see SI text and Fig. S1 for detailed explanation). Thus, in practice the traditional 2-step approach may not allow for an unambiguous interpretation of the replay dynamics. In contrast, our method adopts a more principled approach, namely explicit modeling of drift-diffusion, as a means to systematically disambiguate the speed (drift) and quality (diffusion) of a replay event. Importantly, this approach enables parameter estimation in a single step, allowing a more principled modeling of uncertainty.

### 2.3 Applications to experimental data to investigate open questions about replay

We next apply our modeling framework to investigate the structure and dynamics of hippocampal replay. We will investigate three questions that remain unresolved in research on hippocampal replay. First, does hippocampal replay depict trajectories that evolve at real-world speeds, or is even stationary [25]? Second, does hippocampal replay evolve similarly to Brownian motion (random walk) [31]? Third, how does the structure of the hippocampal “preplay” events (during pre-run sleep) compare with replay events [32, 46]? We analyze a previously published dataset [33] that contains hundreds of simultaneously recorded neurons from the hippocampus when the rats navigated on a one-dimensional linear track (see Fig. 1A,B for an overview of the task paradigm).

As an initial step of the analysis, we fit our full switching dynamics model to neural population response during SWRs collected in [33] (a total of N=16,523 events from 4 rats). A summary of the results is shown in Fig. 4. We find that 12.2% of the events in this dataset can be classified as drift-diffusion dynamics only. In addition, about 6.7% of events are best described as a mixture of drift-diffusion and other types of dynamics. Thus, 18.9% of SWR events contain at least a segment of drift-diffusion period (i.e., at least 20 ms). This fraction is generally consistent with the previously reported fraction of replay events, analyzed using standard Bayesian decoding methods [4, 49]. Unlike prior methods, our method enables a more principled characterization of the rich dynamics within individual events (see examples in Fig. 2B-H and Fig. 6A).

**Figure 4:**
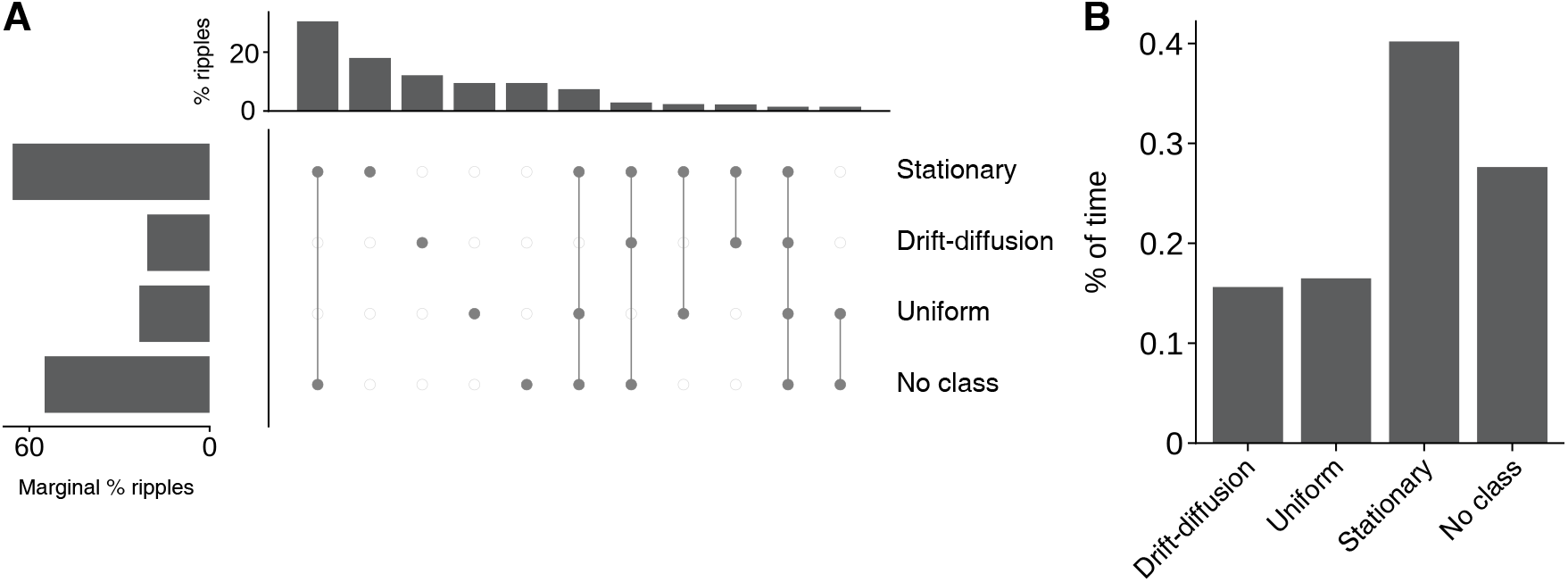
Composition of spatiotemporal dynamics of SWRs. For each SWR, the marginal posterior of dynamics was classified into a time series of regime class {*ĉ*_*t*_} following the convention in Fig. 2B-H. The time series was then partitioned into several segments, each defined as a period in which a single regime class persisted for ≥ 20 ms. **A**. The UpSet plot [48] summarizes the co-occurrence of the segments in all SWRs. Bottom: Dot–line matrix marks which segments are present in each combination. Filled circles denote the presence of a given segment in that combination, connected by lines. Top: Bars show the most frequent segment combinations in SWRs, ranked by the prevalence percentage. Left: Bars show the marginal percentage of SWRs that contain at least one segment of each regime class. **B**. Fraction of total SWR time assigned to each regime class.

### Application 1: Do most replay events exhibit real-world moving speed or even remain stationary?

Recent work suggests that a large fraction of SWR events were either stationary or had a replay speed that is comparable to the animal’s moving speed (*<* 1 m/s) [25]. This result seems to contrast with many previous reports showing that replay is compressed temporally, and its speed is often ten times faster than the animal’s movement speed [3, 7, 49].

In light of this controversy, we next focus on investigating the stationary dynamics of SWRs. While our model fitting results indicate that ∼40% of the time during SWR exhibits stationary dynamics and seem to be consistent with the conclusion of [25], crucially, these results need to be interpreted together with the chance levels, *i*.*e*., the fraction of the stationary dynamics inferred from the shuffled data. We consider three shuffle procedures (Fig. 5). First, time-bin shuffle permutes the spike raster of each SWR across time bins, preserving per-neuron spike counts while disrupting within-event temporal structure. Second, neuron-identity shuffle randomly shuffles spike trains across neurons, preserving the spiking profile of each neuron but breaking the neuron-position mapping. Third, place-field rotation shuffle circularly shifts the place field of each neuron by an independent random offset. We compared the results obtained from real data and three shuffles. While the fractions of stationary events vary slightly across three shuffling methods, the results consistently show that the fraction of hippocampal SWRs with stationary dynamics is higher than the fractions obtained from all three shuffles. However, the differences are relatively small. Specifically, the fraction of SWR events with a substantial stationary period (> 50 ms) is only about 5% more than the shuffles based on all three methods. Together, these results suggest that only a small fraction of SWR events are *bona fide* stationary events, and this fraction is much smaller than that reported in [25].

**Figure 5:**
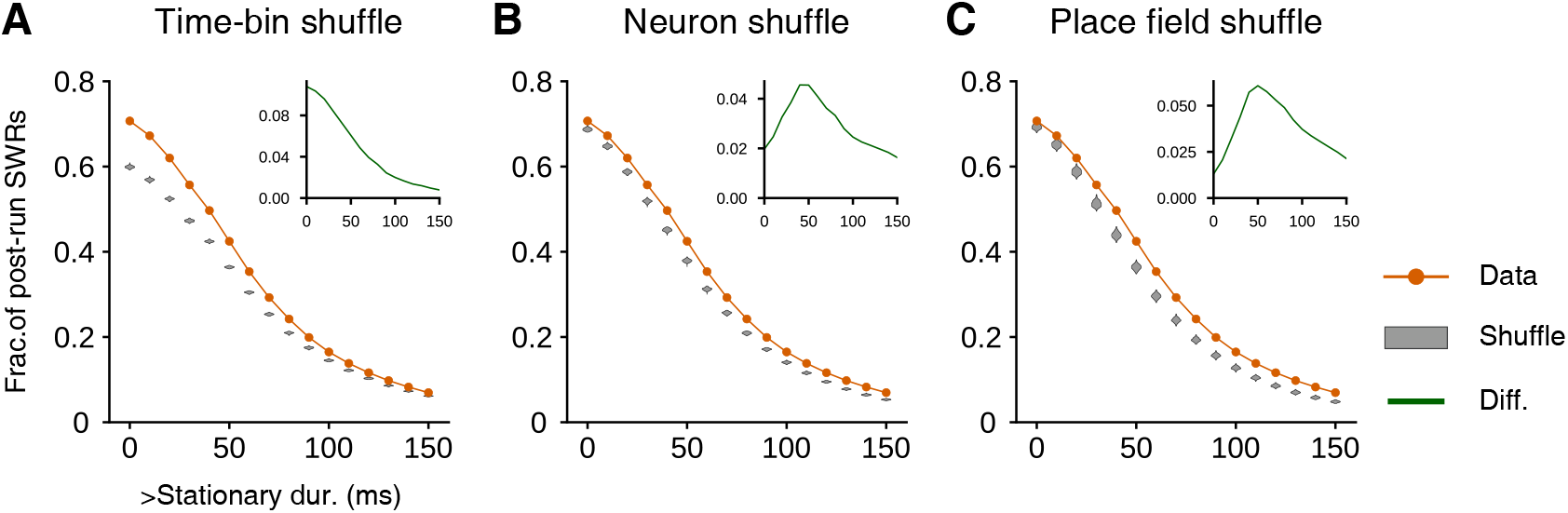
The fraction of stationary events in post-run SWRs slightly exceeds chance levels obtained from three different shuffles. Each plot shows the fraction of SWRs during awake and post-run sleep periods that have at least the minimum stationary duration specified on the x-axis. Orange lines indicate the fraction of SWRs in data. Gray violins indicate the distribution of fractions from 120 shuffles per SWR. Inset shows the difference across thresholds between fraction in data and mean fraction in shuffle. Positive value indicates a higher fraction than expected under shuffle and negative value indicates lower. Columns correspond to three shuffle controls: time-bin (**A**), neuron-identity (**B**), and place-field rotation (**C**).

Next, we examine a subset of SWR events with marginal posterior of drift-diffusion dynamics that exceed 0.7 for at least 50 ms. These events are particularly interesting because they exhibit strong sequential structures and thus are consistent with the classic notion of replay events. These selected events account for 17.3% of the total events (N=2861 events). Fig. 6A shows example SWR events with varying replay speed. The distribution of the replay speed inferred for these events is shown in Fig. 6D, with a median of 4 m/s. The speed for these replay events is substantially higher than that of running (Fig. 6B) or movement periods (Fig. 6C). These results are insensitive to the exact criteria used to select these events. Thus, for the subset of SWR events that exhibit substantial drift-diffusion dynamics, the speed of the drift is one order of magnitude faster than the running speed, generally consistent with earlier reports on replay [8, 49].

**Figure 6:**
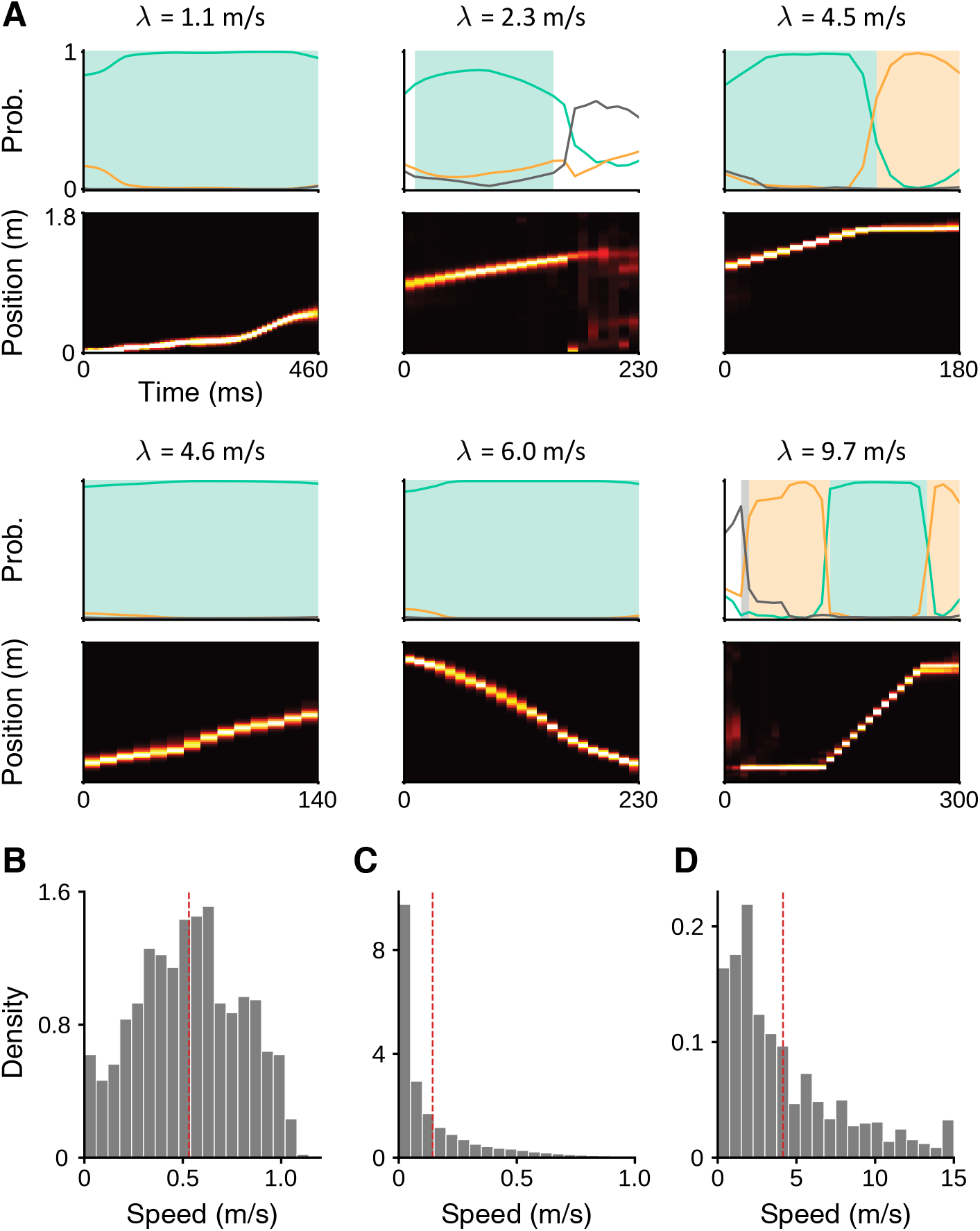
Speed distributions for running, behavior, and replay events. **A.** Example SWR events exhibiting different speeds. Plotting conventions follow Fig. 2B-H. **B**. Histogram of per-lap running speed, estimated by fitting a linear regression to the running trajectory of each lap on the linear track. **C**. Histogram of instantaneous behavioral speed, estimated using a finite-difference derivative of positions during running. **D**. Histogram of replay speed, estimated using the inferred drift rate *λ*. Only drift-diffusion events, namely SWRs whose marginal posterior of drift-diffusion regime exceeded 0.7 for at least 50 ms, are considered in **D**. The red dashed lines in **B-D** indicate the mean of each speed distribution: run, 0.53 m/s; behavior, 0.14 m/s; replay, 4.1 m/s.

Our results suggest that the subset of SWR events (generally 10%-20% of all SWR events) with high substantial sequential structure indeed exhibit a speed that is approximately an order of magnitude faster than the animal’s typical running speed, consistent with classic studies of replay [8, 49]. In addition, our results provided evidence for the existence stationary dynamics in some SWR events [25]. However, by carefully considering the chance levels from the shuffled data, we find that stationary events only account for a small fraction of SWRs.

### Application 2: Does hippocampal replay in 1-D environment resemble Brownian motion?

Although earlier studies of replay emphasize the linear-sequential structure of replays, we found that the decoded trajectories of many SWR events have substantial variability compared to the path of the animals’ behavioral trajectories. Intriguingly, a recent study [31] on hippocampal activity in a 2-D open arena argues that replays resemble Brownian motion (but see [26]). Though this a conceptually provocative result, it remains unclear whether similar results hold in other datasets and experimental settings, particularly that of 1-D environments, where replays are typically studied.

One of our results reported above seems to contradict the proposition that Brownian motion generally describes the dynamics of SWRs, that is, a considerable proportion of SWRs exhibit a substantial drift (Fig. 6B). However, it remains possible that the noise may cause the empirically estimated drift to substantially exceed zero even when sampling the trajectories from a pure diffusion process. To rule out this possibility, we generated a null distribution of the estimated speed from the diffusion process, by sampling and analyzing trajectories from diffusion models. We find that this null distribution is substantially different from the distribution obtained from the rat data (Fig. S3). In particular, the speeds obtained from the simulated diffusion process are substantially slower. These results suggest that the subset of drift-diffusion events identified by our method are unlikely to be characteristically generated by Brownian motion.

To make the results directly comparable to prior work, we further analyzed the data using an approach generalized from [31], which examines how the root-mean-square displacement (RMSD) scales with the time interval (Fig. 7A). Their key idea is that the Brownian motion predicts that RMSD should increase linearly with the square root of time interval 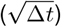, i.e., RMSD 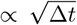. Empirically, Stella et al. [31] reported that the slope between these two quantities is reasonably close to 0.5 in a log-log plot. To evaluate this prediction independently, we generalize the previous analysis framework to more general drift-diffusion dynamics. Importantly, if the replay trajectory follows a drift-diffusion process, the mean-squared displacement (MSD) should be characterized by a quadratic rather than linear function, that is, MSD = (*λ*Δ*t*)^2^ + *σ*^2^Δ*t*. Under this view, Brownian motion is only expressed in the special case of zero drift, in which MSD = *σ*^2^Δ*t*.

**Figure 7:**
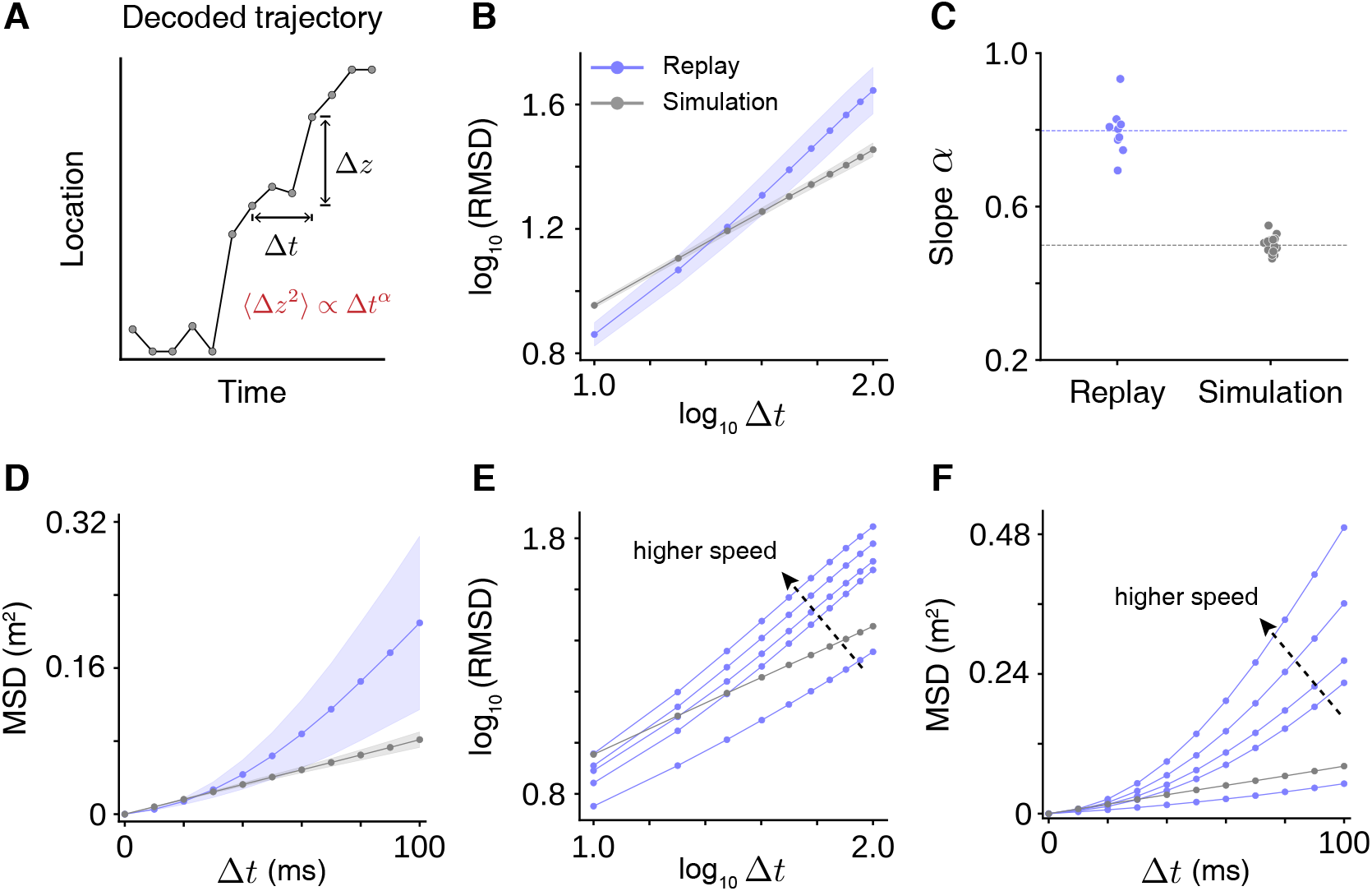
Replay trajectories differ from pure diffusion dynamics (Brownian motion), and are more consistent with drift-diffusion. **A.** Schematic diagram of the calculation of position displacement Δ*z*, mean-square displacement (MSD) ⟨Δ*z*^2^⟩, and root-mean-square displacement (RMSD) 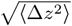 for the time interval Δ*t* from a replay trajectory. **B**. The log-log relationship between RMSD and Δ*t* for replay (purple) and simulated (gray) trajectories. Note that replay trajectories were inferred under a fixed pure diffusion model 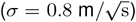. Simulated replay trajectories were generated from the same pure diffusion model [26]. Solid lines and shaded bands indicate the mean and *±*1 standard deviation across sessions, respectively. **C**. Slopes of linear regression fit to log(RMSD)-log(Δ*t*) data shown in **B**. Each dot represents an estimated slope from one session. Dashed horizontal lines indicate the mean across sessions: 0.79 for replay and 0.50 for simulation. **D**. The relationship between MSD and Δ*t* for replay (purple) and simulated (gray) trajectories. Solid lines and shaded bands indicate the mean and *±*1 standard deviation across sessions. **E**. The log-log relationship between RMSD and Δ*t* for replays with different estimated speed (purple) and simulations (gray). Replay events from all sessions were grouped based on five speed levels: *λ* ∈ [0, 2), [2, 4), [4, 6), [6, 8) and [8, 10] m/s. The slopes of linear regression fits are 0.61, 0.85, 0.84, 0.88 and 0.90, respectively. Gray lines show the same analysis using simulated trajectories from the pure diffusion model, where the slope is 0.50. **F**. The relationship between MSD and Δ*t* for replays with different estimated speed (purple) and simulations (gray). The definition of speed groups is identical to **E**. Gray lines show the same analysis using simulated trajectories from the pure diffusion model.

In examining log-log plots of RMSD vs. Δ*t* (limited to drift-diffusion events, namely events with drift-diffusion time lasting at least 50 ms), we observed a slope of approximately 0.8, substantially greater than 0.5. The results are consistent across nine sessions (mean=0.80, std=0.06) and deviate substantially from the predictions of Brownian motion (Fig. 7B,C). Importantly, to avoid biases toward finding systematic drifts, we used a fixed diffusion model (with 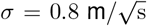, zero drift) to decode the trajectories [26].

Our theoretical analysis reveals one important limitation of the approach based on the slope between RMSD and time interval in the log-log space. The issue is that, while the slope in the log-log space may be adequate to recover the exponent *α* for a power-law relation MSD = *C*(Δ*t*)^*α*^, it is insufficient to test the more general prediction of drift-diffusion models in the form of MSD = (*λ*Δ*t*)^2^ + *σ*^2^Δ*t*. To overcome this limitation, we further examined how MSD scales with time interval in the original scale for drift-diffusion events. Consistent with the prediction of the drift-diffusion dynamics, we find that the relationship between MSD and time interval can be well described by a quadratic, rather than a linear, relationship (Fig. 7D). A further prediction of the drift-diffusion model is that the relationship between MSD and time interval should be systematically modulated by the drift rate. To test this, we sorted and separately analyzed the SWR events with different drift rates, finding an even stronger quadratic relationship for the events with higher speed (Fig. 7F). In the log-log plots, this yielded a slope that was closer to 1 for high speed events (Fig. 7E). These results provide strong empirical support for the drift-diffusion dynamics of replays.

A key advantage of our modeling approach is that it potentially enables the identification and characterization of different types of replay dynamics. Indeed, our empirical results in Fig. 4 suggest that SWRs exhibit different types of dynamics. If these different types of events are indeed meaningful, one substantive prediction, generalizing the one suggested specifically for Brownian motion above, would be systematically different scaling relationships between MSD and time interval. To test this prediction, we additionally analyzed the subset of events that contain only stationary dynamics, as well as the subset of events that contain only uniform dynamics. Consistent with our prediction, for SWRs with stationary dynamics (Fig. S5A-C), the slope in the log-log plot is substantially smaller than 0.5 (mean=0.26, std=0.13). Furthermore, for events with uniform dynamics (Fig. S5D-F), the slope is also smaller than 0.5 (mean=0.39, std=0.07). When all SWR events are combined, the resulting slope between RMSD and time interval in the log-log plots is 0.61, which is closer to, yet remains different from, the prediction of Brownian motion (Fig. S5G-I). Thus, these results strongly suggest that the scaling relationship between MSD and time is heterogeneous in SWR events with different types of dynamics.

### Application 3: Is there evidence for preplay during SWRs?

It has been suggested that the hippocampal “preplay” sequences of neural activity before the animal has any experience in a given environment [32, 50]. While preplay potentially has important implications for the mechanisms and functions of hippocampus, the existence and prevalence of preplay remain controversial [46]. This is in part due to the difficulty of precisely characterizing the dynamics of the SWR events. Using our method, we sought to re-examine the evidence for preplay, and if preplay exists, how its dynamics compare with replay dynamics after spatial experience in a given environment.

We analyzed the SWRs during pre-run sleep (data from ref [33], a total of N=6,749 events from 4 rats). We first quantified the composition of dynamics (see Fig. S6). For pre-run sleep, drift-diffusion dynamics accounted for 5.8% of total SWR time, which is substantially lower than in post-run sleep (15.6%). At the event level, 5.6% of pre-run SWRs can be classified as drift-diffusion dynamics only and 6.3% contained at least one drift-diffusion segment. These numbers are again much smaller than those from post-run (12.2% and 18.9%, respectively). Thus, when aggregated across events, pre-run SWRs exhibit less sequential structures than post-run SWRs.

To assess whether the observed fraction of preplay events is significantly higher than chance levels, we conducted three shuffle analyses mentioned earlier (time-bin shuffle, neuron-identify shuffle, and place-field rotation shuffle) on preplay using data from pre-run sleep periods [46]. We further compare the results with those obtained from the shuffle analyses on post-run sleep SWRs. Taking advantage of our drift-diffusion framework, we used the duration of drift-diffusion dynamics within an event as a metric to characterize the sequentiality of an event (Fig. 8A-F). Across a range of thresholds, we computed the fraction of SWR events that exceeded each threshold to assess statistical significance. We find that the results diverge across the three shuffle methods. Based on either neuron-identity shuffle or place-field rotation shuffle, we did not find evidence for preplay, consistent with results reported in [46]. However, for time-bin shuffle, the fraction of events that exhibited sequentiality was higher than the chance level, consistent with [50]. These results may help reconcile the previous discrepancy on the existence of preplay from studies that used different shuffle procedures [46, 50]. It is also interesting to note that the time-bin shuffle has been suggested to be relatively less conservative than the neuron-identity and place-field shuffles [20], which also underscores the finding of significant preplay structure as surprising.

**Figure 8:**
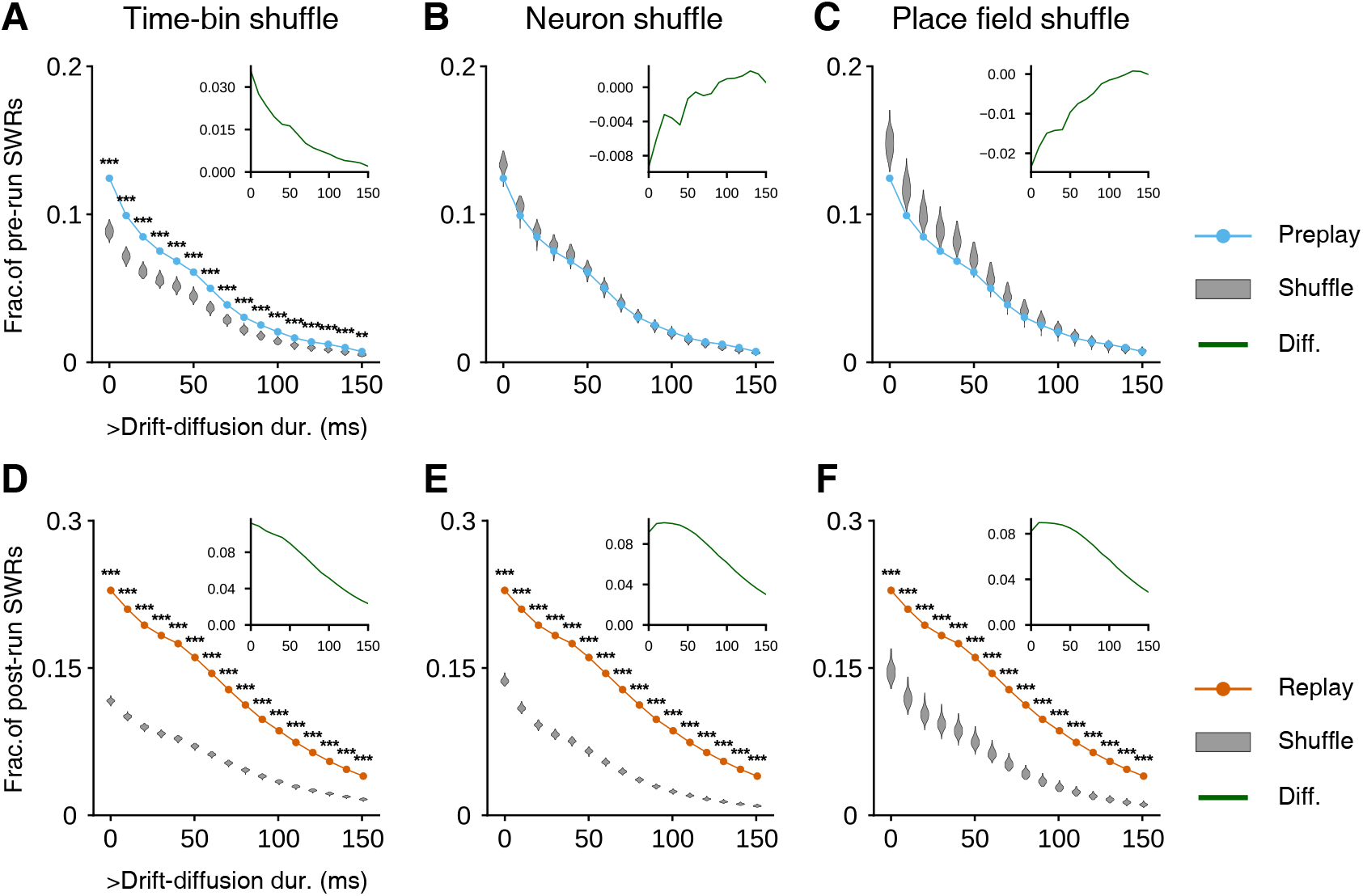
Preplay events show substantially weaker temporal structure than replay. **A-F**. Each plot shows the fraction of SWRs that have at least the minimum drift-diffusion duration specified on the x-axis. Colored lines indicate the fraction of SWRs in data. Gray violins indicate the distribution of fractions from 120 shuffles per SWR. Asterisks mark the significance of the fraction in data being higher than the shuffle (one-sided permutation test, *** *p <* 0.001 and * *p <* 0.05). Inset shows the difference across thresholds between fraction in data and mean fraction in shuffle. Positive value indicates a higher fraction than expected under shuffle and negative value indicates lower. Rows correspond to two subsets of SWRs: pre-run SWRs (**A, B, C**) and post-run SWRs (**D, E, F**). Columns correspond to three shuffle controls: time-bin (**A, D**), neuron-identity (**B, E**), and place-field rotation (**C, F**).

When comparing pre-run and post-run SWRs, we consistently find that, across the entire threshold range, the fraction of events exceeding any given threshold was higher for post-run than pre-run SWRs. This suggests that at the level of individual events, pre-run SWRs also exhibited shorter drift-diffusion segments than post-run SWRs, mirroring the pooled result in Fig. 4. We also compared the distributions of drift-diffusion durations against their corresponding null distributions based on shuffle (Fig. S7). Post-run SWRs showed differences from all three controls with strong significance (p < 0.001 for each; one-sided permutation test). In contrast, pre-run SWRs differed only from the time-bin shuffle (p < 0.001; one-sided permutation test) and not significantly from the neuron-identity or place-field rotation shuffles. These results suggest that replay after experience can be detected robustly, while preplay is detectable only when neuron-position mappings are preserved, as in time-bin shuffle, but not in neuron-identity and place-field shuffles.

The results above suggest that hippocampal population activity during pre-run SWRs is coordinated between potential (unexperienced) spatial locations at a level that is higher than chance. We next examined whether such coordination happens at a timescale that is comparable to the classic replay events (typically around 100 ms). By taking the difference of the distribution of drift-diffusion durations between real data and shuffles (Fig. 8A-F inset), we obtained an estimate of the fraction of *bona fide* drift-diffusion events with relatively long durations. According to this metric, for post-run SWRs, the fraction of events that exhibit drift-diffusion longer than 100 ms is about 5.16% according to time-bin shuffle. For preplay, this fraction is 0.64%. Thus, hippocampal population activity during pre-run SWRs exhibits slightly higher sequentiality than time-bin shuffled data, yet long temporal sequences are particularly rare.

Together, these results provide evidence for pre-existing structures in place-cell population activity before rats have spatial experience in a given environment. However, the level of coordination in these structures is substantially weaker than that seen in replay. In particular, we found that long sequences during pre-run SWRs are rare, much weaker than replay sequences after spatial experience.

## 3 Discussion

We have developed a new framework, conceptually and statistically, for analyzing and understanding the dynamics of replays. This framework is formulated based on drift-diffusion dynamics while allowing the switching between several types of dynamics. Drift-diffusion models have been widely used to model the neural basis of decision-making [51–53]. Here we show that the drift-diffusion framework also provides a powerful framework to study the dynamics of the hippocampus. Compared to standard methods for analyzing replays, there are several important differences that make our method advantageous. First, using a generative modeling approach, our method enables principled probabilistic inference on the structures of replays that fully takes into account the uncertainty in estimation. Second, our framework explicitly models speed and quality of classic replay events (i.e., drift-diffusion events) with drift and diffusion parameters. Third, with the capability of switching between different types of dynamics, our method is able to characterize the putatively rich dynamics in a broader set of hippocampal SWRs, which continues to remain an important open question. Evaluation and comparison of our method to others reveal that our method allows for more direct and unambiguous interpretation of replay dynamics.

Application of our modeling framework to a large-scale hippocampal dataset revealed several insights that helped resolve controversies about replays. First, our results clarify the recent debate on the replay speed and stationary dynamics during SWRs. Consistent with the results reported in [25], we find that some SWR events contain segments of stationary dynamics. However, the fraction of *bona fide* stationary events is likely much smaller than that reported in [25], when considering the chance levels obtained from the shuffled data. Somewhat surprisingly, a large fraction of events from the shuffle data exhibit stationary dynamics. This suggests that stationary dynamics observed in decoding may arise from multiple sources and by itself cannot be interpreted as conclusively evidence for strong coordination of neural population activity. While the exact reasons for observing many stationary events in the shuffle data remains to be determined in future work, we hypothesize that the following two factors may contribute it. First, the biases in the decoded posterior distribution due to the heterogeneity of the sampled place cell code in the experiment may cause segments posterior to be consistent over space. Second, if a small subset of neurons consistently exhibit high firing rates during segments of SWRs, stationary dynamics may emerge. We also find that, for a subset of SWR events that exhibit consistent sequential structures, the speed is about 10 times faster than the animal’s running speed. These events account for 10-20% of the SWRs, which is consistent with the classic replay sequences during SWRs. Thus, our results shed light on the discrepancy of the results in [25] and classic studies of replay. Our results also highlight the rich and diverse types of neural dynamics during SWRs. The role that different types of neural dynamics play in learning and memory remains unclear–an important question that needs to be determined in future work.

Second, our results clarify the dynamics of replay events, at least in 1-D linear tracks. By carefully analyzing the scaling relationship between the MSD and time interval, we found that the two follow a quadratic relationship for events with high sequentiality. This result is highly consistent with the prediction of drift-diffusion dynamics, but not with the Brownian motion as proposed in [31]. SWR events with higher speed lead to a more substantial divergence from Brownian motion. We also found that different types of events exhibited different scaling relationships. For stationary events and uniform events, the relationship is sublinear. These results provide further support for the distinctness of different types of dynamics identified using our method. Our results suggest that it is more informative to analyze the scaling relationship between MSD and time interval in the original scale, rather than the log-log scale proposed previously [31]. The latter may confound potential contributions from diffusion and drift. While our results are based on data collected in 1-D environments, they motivate revisiting or additional exploration of replay dynamics in 2-D, or in additional environment and behavioral settings. In particular, it would be interesting to examine or re-examine the 2-D data to see whether a general quadratic relation also holds in 2-D for the replay events they identified. It would also be important to examine whether there is substantial heterogeneity in the dynamics of SWR events in 2-D. One possibility is that, for SWRs in 2-D environment, the dynamics are also heterogeneous as in 1-D, yet the fractions of SWR events with different types of dynamics differ from those in 1-D. When combining all events together, this may result in an overall slope close to 0.5 in the log-log scale. Regardless, our results clearly demonstrate that the dynamics of SWRs are diverse, and cannot be adequately described by Brownian motion in 1-D environments.

Third, we find evidence supporting the existence of weak pre-existing temporal structures before spatial experience in a given environment, consistent with the idea of preplay [32]. However, these pre-existing structures appear to be detected with particular analysis approaches–in particular, we found population structure beyond randomization only through time-based shuffle. Furthermore, our results reveal that long drift-diffusion sequences that are comparable to classic replay events with a timescale of 100 ms are exceptionally rare (less than 1% of SWRs in pre-run sleep). Even with these results, caution is in order, as time-based shuffle may eliminate temporal structure in individual neurons, thus it may provide a weaker null distribution. Furthermore, given that the animals may have been exposed to environments with somewhat similar spatial structure, sequences observed during pre-run SWRs may still reflect a type of replay of analogous experiences [9, 54, 55]. Exactly what properties of the pre-existing hippocampal circuits lead to the higher-than-chance sequential structure during pre-run sleep remains to be determined [56]. One possibility is that the structure during pre-run sleep is caused by the cellular correlation between “rigid” place cells that is preserved across environments [57].

Earlier studies have used state-space models to decode positions and reconstruct “mental” trajectories from hippocampal activity during running and sleep, although they did not explicitly characterize the spatiotemporal structure of replay [28–30, 38, 39]. More recently, state-space models have been applied to explicitly model dynamics during SWRs [25, 26]. In particular, [25] proposed a method for analyzing hippocampal replay by considering switching dynamics during decoding. There are two crucial differences between the method in [25] and the method presented here. First, the parameters of their method were manually specified rather than inferred from data. Second, their method does not model drift, in part as a consequence of the manual specification. Instead, it relies solely on Brownian motion as its spatially “continuous” dynamics to capture the sequential structure of replay events. As we have shown, Brownian motion is inadequate for capturing sequential replay dynamics. These limitations may introduce biases in estimating dynamics and cause decoding errors in estimating positions. For example, the replay speeds based on their method could be underestimated if the replay in neural data has a substantial drift, which does not conform to the manually specified structure in the model. In contrast, our method directly estimates two essential model parameters from individual replay events and is thereby more flexible. Another recent study [26] formulated HMM models to study transition dynamics in 2-D open-field environments. They found that the momentum of the trajectory is important for fitting replay events. One important difference is that their model does not allow for switching between different types of dynamics. Also, their model is a second-order HMM, our model is first-order. From what we have shown, a first-order switching HMM appears sufficient to capture the key dynamical patterns of replay. Moreover, its dynamics and parameters (e.g., drift and diffusion) map directly onto first-order transition statistics, leading to a more transparent interpretation of how replay trajectories evolve over time. Our first-order formulation also simplifies the probabilistic inference procedure and potentially improves parameter identifiability, leading to more stable and accurate characterizations of replay dynamics.

Extensions and additional applications of the proposed modeling framework are possible. First, our current implementation primarily deals with 1-D track or circular environment, two popular environments that have been used to study hippocampal replay. Other types of environments have also been used in studying replay, such as 2-D open-field arenas and 1-D environments with more complex geometry. Adaptation of our method to deal with these environments would require modifications to the probabilistic transition matrix for latent positions. One simple generalization of our method to 2-D environments would be to assume that dynamics in 2-D factorize into two individual dimensions, and apply our method to each dimension separately. Second, another natural extension would be a switching drift-diffusion model that switches between multiple drift-diffusion processes with distinct parameters. However, in practice, the limited data per SWR event may prevent one from estimating the parameters reliably. In this paper, we therefore adopt a more parsimonious model by explicitly including two stereotyped dynamics (i.e., stationary and uniform sampling) alongside a single drift-diffusion component. This choice preserves tractability while maintaining flexibility to capture the spatiotemporal heterogeneity of individual SWRs. Crucially, by allowing stationary and uniform regimes in the model, the method can isolate SWR segments with substantial drift and spatial coherence. Thus, the duration of drift-diffusion segment for individual events provides a measure of the strength of sequential replay. Third, while we only focus on replay sequences during SWRs, our method is in principle applicable to the study of theta sequences in rodents during running [58, 59]. Theta sequences represent another important type of dynamical structure in the hippocampus [60–62] and entorhinal cortex [63], which may have equally important functional implications for the role of hippocampus or other areas in cognition. Previous analysis typically relied on analyzing the averaged slope of the theta sequences. Using our method, one may potentially reveal the sequentiality and variability of the dynamics within individual theta sequences.

A better understanding of replay dynamics in the brain may help inform the development of better learning algorithms in artificial intelligence. Algorithms based on replaying past experiences have been developed and shown to improve performance across a variety of tasks in reinforcement learning [65, 66]. Yet most current AI algorithms still struggle in continual learning without catastrophic forgetting [67–69]. Developing brain-inspired algorithms to replay past experiences efficiently [70, 71], or to sample from a generative model built on past experiences (i.e., generative replay) [72–74], remains a promising path forward [75, 76]. More accurate and interpretable data-driven techniques, such as the one presented here, will likely be instrumental toward clarifying the exact role of replay in natural cognition, thereby informing brain-inspired replay mechanisms for continual learning in AI.

Hippocampal SWRs have direct implications in cognitive functions. It was reported that disrupting SWRs in rodents impairs spatial memory in an AD-developmental mouse model [77] or spatial learning in wild-type [78], whereas enhancing SWRs may improve subsequent task performance [79]. Recent work shows that in a mouse model of Alzheimer’s disease, the reactivation of hippocampal activity during SWRs was disrupted [80]. Most prior work used simple metrics such as the frequency of SWRs. Using our method to analyze the detailed dynamics during different task conditions may reveal new insights into the relationship between the properties of SWRs and behavior. Finally, adapting our method to the online setting may enable the design of more targeted, dynamics-specific, SWR-based neurofeedback protocols for cognitive perturbation and enhancement [81, 82].

## Acknowledgments

We would like to thank Brad Pfeiffer and David Foster for generously sharing their data. We thank Laura Colgin and Kenneth Kay for helpful discussions and their comments on an earlier version of the paper. This research is supported by NSF 2318065, and a Sloan Research Fellowship to X.-X. Wei. The funders had no role in designing and conducting this research, decision to publish, or preparation of the manuscript.

## 4 Methods

### 4.1 Description of the datasets

We used electrophysiological recordings in Pfeiffer and Foster (2015) [33]. In these experiments, four adult Long-Evans rats (Janni, Harpy, Imp and Ettin) performed back-and-forth runs on a one-dimensional linear track. A total of nine recording sessions were analyzed: four sessions from Janni, two from Harpy, one from Imp, and two from Ettin. Seven of these sessions followed the standard three-epoch structure: pre-run sleep in an isolated box, active running on the track, and post-run sleep in the same box. In the remaining two sessions, after an initial pre-run sleep epoch, the animals underwent three alternating run and post-run sleep epochs. Each run epoch lasted for approximately 40 min, and each pre- or post-run sleep epoch lasted for 40-60 min. The linear track was 1.8 m in length and 6.35 cm in width, expanding to 12.7 cm at each end, and was enclosed by 3.8 cm high walls. At both ends of the track, reward wells were filled out of view with 200–250 *µL* of chocolate-flavored liquid to encourage traversal. During each session, dorsal CA1 place cells were recorded as the animals navigated the track. Animal position and head direction were simultaneously monitored at 60 Hz using an overhead camera. More details about the experiments can be found in [33].

### 4.2 Place field estimation

Place fields were estimated using the neural responses while the animal was moving at speeds larger than 5 cm/s. The linear track was discretized into 2 cm spatial bins. For each neuron *i*, the neural activity vector was constructed by counting the number of spikes that occurred within each spatial bin. This vector was then divided by a spatial occupancy, computed by accumulating the time the animal spent in each bin. The resulting firing rate was smoothed using a Gaussian kernel with a standard deviation of 5 cm. The smoothed firing rate vector, denoted *f*_*i*_(*z*), was taken as the place field of neuron *i*. A neuron was classified as a place cell if the peak firing rate of its place field exceeded 1 spike/s. Only neurons identified as place cells were included in subsequent analysis.

### 4.3 Sharp-wave ripple detection

Sharp-wave ripples (SWRs) were detected following the procedure of Karlsson and Frank (2009) [9]. Raw local field potential (LFP) data were first band-pass filtered between 150 and 250 Hz to isolate the ripple-band oscillations. The instantaneous amplitude of the band signal was then obtained by taking the absolute value of its Hilbert transform and smoothing with a Gaussian kernel (*σ* = 4 ms). Candidate ripple events were identified as periods during which the smoothed amplitude exceeded the session-wide mean by three standard deviations. Each event’s start and end times were extended backward and forward, respectively, to the time when the amplitude first crossed above and then fell below the mean. Overlapping detections across tetrodes or channels were merged into single events. Ripple event boundaries were further refined by excluding edge time bins with no spiking activity. Only events lasting no less than 50 ms were retained.

### 4.4 State-space modeling

#### 4.4.1 Transition dynamics

Temporal transitions of replayed positions were modeled with several categories of dynamics, including drift-diffusion, stationary and uniform processes.

##### Drift-diffusion

We start from the continuous-time differential equation for drift-diffusion process

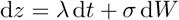

where *λ* and *σ* are the drift and diffusion parameters, respectively, and *W* is a Wiener process with 𝔼[d*W*] = 0 and Var[d*W*] = d*t*. Discretizing this stochastic differential equation with time-bin width d*t* gives a Gaussian transition kernel,

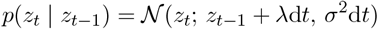

where 𝒩 (*z*; *µ, σ*^2^) denotes the normal density of *z* with mean *µ* and variance *σ*^2^.

##### Stationary

Similarly, we write the continuous-time differential equation for stationary dynamics

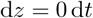

meaning the latent position is held fixed in time. Discretizing with bin width d*t* yields the trivial transition kernel:

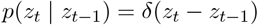

where *δ*(·) is the Dirac delta.

##### Uniform

Here we assume complete absence of temporal structure. Each latent state is resampled independently at every time step. There is no meaningful continuous-time limit, so in discrete time

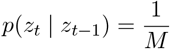

Thus, every next state is equally likely, irrespective of the previous state.

Under certain parameter limits, the drift-diffusion dynamics reduces to other two candidate dynamics. In the limit *λ* → 0 and *σ* → 0, the Gaussian transition collapses to a point-mass at the previous state, reproducing the stationary model. As *σ* → ∞, the Gaussian becomes flat over the track, and the transition matrix approaches the uniform model. In practice, during SWR events dominated by a single dynamical regime, these extreme parameter settings lead to near-equal marginal posterior probabilities for the drift-diffusion versus stationary or uniform categories.

#### 4.4.2 Switching HMMs

An individual SWR event may switch between multiple regimes rather than following a single type of dynamics [25]. We developed a latent-position HMM model based on a two-level Markov-switching framework. The full joint distribution in this model can be described as

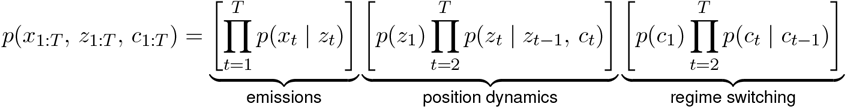

We model the emission probability *p*(*x*_1:*T*_ | *z*_1:*T*_) as conditionally independent Poisson observations given the latent location. Writing *x*_*t,i*_ for the spike count of neuron *i* at step *t*, and *f*_*i*_(*z*) for its place field at location *z*, the emission likelihood factorizes as

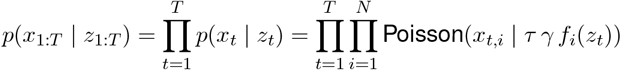

where

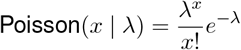

Here *τ* = 10 ms is the bin width, and *γ* is a global gain factor that accounts for the increased firing rates during SWR events relative to locomotion.

To allow individual SWR events to exhibit mixtures of our three candidate dynamics, we augment the latent-position HMM with a discrete regime chain *c*_*t*_ ∈ {1, 2, 3}, where states 1, 2 and 3 index drift-diffusion, stationary and uniform dynamics, respectively. We write

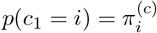

where 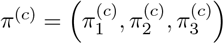 is the initial distribution over regimes, and

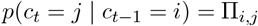

where Π is the regime-transition matrix. We further define the transition matrix as

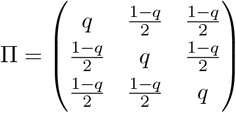

This formulation assumes equal self-transition probability *q* across all regimes and uniform switching probability between regimes. The parameter *q* is learned from the data on an event-by-event basis.

Given the current dynamical regime *c*_*t*_, the latent position *z*_*t*_ then evolves according to the corresponding transition probability kernel:

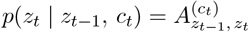

with *A*^(1)^, *A*^(2)^, and *A*^(3)^ defined in the above section for drift-diffusion, stationary and uniform dynamics. The joint probability over (*c*_1:*T*_, *z*_1:*T*_) thus factorizes as

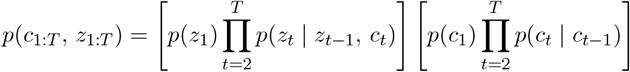

##### Optimization of the model

For each SWR, we fit the three free parameters

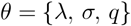

by maximizing the marginal likelihood of the observed spikes under the switching HMM,

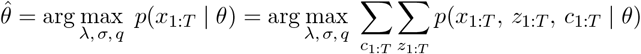

We compute *p*(*x*_1:*T*_ | *θ*) via a forward pass over the two-level Markov chain (*c*_*t*_, *z*_*t*_). We define the forward variable

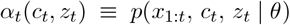

which satisfies the following initialization at *t* = 1:

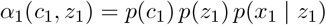

and recursion (*t* = 2, …, *T*):

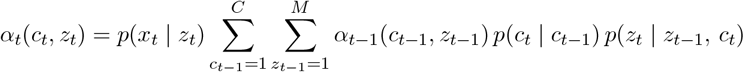

Finally, the marginal likelihood is the total mass of the forward variable at *t* = *T* :

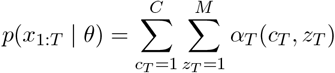

In practice the computation happens in the log scale, thus yielding the overall log-likelihood. The negative log-likelihood serves as the loss function for optimization. Optimization is implemented with `scipy.optimize.minimize` using the L-BFGS-B algorithm, enforcing the box constraints

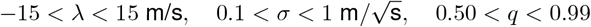

We use default convergence criteria provided by SciPy for termination: the relative objective decrease ≤ 2.2 *×* 10^−9^, or the projected-gradient *ℓ*_∞_-norm ≤ 10^−5^, or upon reaching the iteration limits 15000.

##### Inference

We compute the joint posterior over regimes and positions at each time *t*,

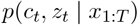

via a standard forward–backward pass. The forward recursion (as discussed in optimization) yields

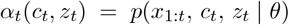

and the backward recursion defines

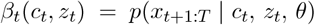

with

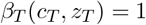

and for *t* = *T* − 1, …, 1:

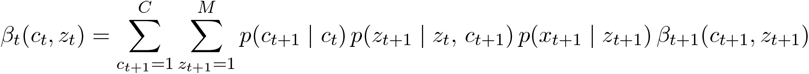

The smoothed joint distribution then can be computed as

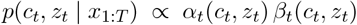

Marginal posteriors can be obtained by marginalization:

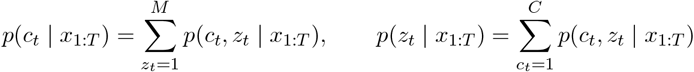

from which we extract both a point estimate of the replayed trajectory and a segmentation into dynamical regimes.

##### Simulation

Our switching HMM is a generative model. We can simulate both latent regimes, latent positions, and spike-count observations given ground-truth parameters

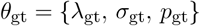

and place fields {*f*_*i*_(*z*)}. For a sequence of length *T*, we proceed as follows: Initialize:

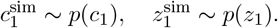

Latent-state evolution (*t* = 2, …, *T*):

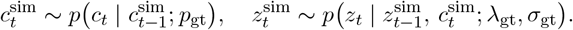

Observation sampling:

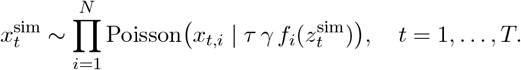

This yields synthetic spike-count data 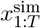 together with their true latent trajectories 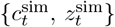. We then can apply the same forward–backward inference and gradient-based parameter optimization to verify the ability to recover *θ*_gt_.

### 4.5 Classic Bayesian decoding and replay metrics

We next describe the procedures for applying standard Bayesian decoding to study replays. SWRs were first binned into non-overlapping 10 ms windows, giving spike-count vector **x**_*t*_ = (*x*_1_(*t*), …, *x*_*N*_ (*t*)). At time *t*, we computed the posterior over *M* spatial bins as

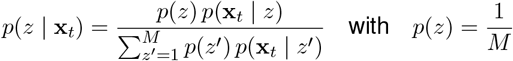

The likelihood was modeled as independent Poisson spiking across cells:

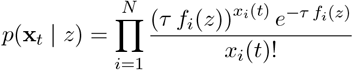

where *τ* = 0.01 s is the time-bin width and *f*_*i*_(*z*) is the place fields of neuron *i* at location *z*. The denominator ensures ∑_*z*_ *p*(*z* | **x**_*t*_) = 1.

We then extracted a decoded trajectory by taking the maximum-a-posteriori estimate at each *t*,

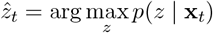

and concatenating 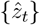 across the SWR.

#### Absolute correlation

We computed the absolute weighted Pearson correlation between time bin index *t* and position bin *z*_*k*_ under the posterior *p*(*z*_*k*_ | **x**_*t*_). We first compute the weighted means

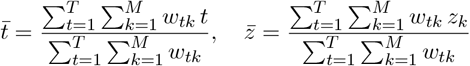

Next, we form covariance

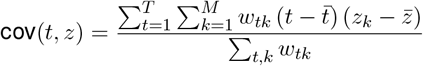

and weighted variances

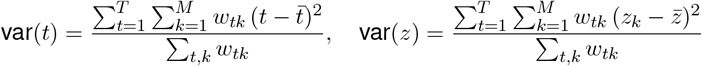

The Pearson correlation is then

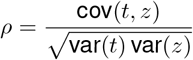

and we report the absolute correlation

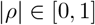

where values near 1 indicate a sequential trajectory and values near 0 indicate little or no monotonic relationship between decoded position and time.

#### Max jump distance

We computed the maximum spatial jump between two time bins within each SWR. From the MAP trajectory 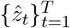, we first formed the sequence of jump magnitudes 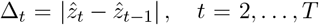, *t* = 2, …, *T*. The max jump distance is then defined as *D*_max_ = max_2≤*t*≤*T*_ Δ_*t*_. High values of *D*_max_ indicate large noise in the decoded trajectory, whereas low values reflect more spatially continuous sequences.

#### Replay speed

We fit a simple linear regression to the MAP trajectory 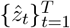 as a function of time bin index *t*: 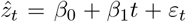. We then define replay speed as the absolute value of the fitted slope |*β*_1_|, which has units of spatial bins per time bin. To express this in physical units (e.g., cm/s), we multiply it by the spatial bin size and divide it by the time bin width.

### 4.6 Synthetic data experiments

#### Parameter identifiability

We ran two simulation experiments to assess the identifiability of drift-diffusion parameters and the accuracy of regime sequence recovery in our switching HMM. In both experiments, we constructed a synthetic population of 300 place cells whose place fields are defined as Gaussian bumps 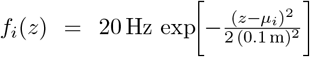. The preferred positions *µ*_i_ uniformly tile a 4 m one-dimensional linear track.

##### Drift-diffusion regime

To test recovery of drift-diffusion parameter, we defined a partial 2-D grid of ground-truth drift and diffusion values. We swept 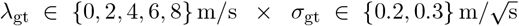 and 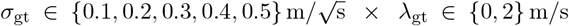. For each (*λ*_gt_, *σ*_gt_) pair on the grid, we simulated 100 independent SWRs as follows. First, generate a 300 ms latent trajectory {*z*_*t*_} by evolving the drift-diffusion process with specified *λ*_gt_ and *σ*_gt_. Next, use the place-cell population to sample Poisson spike counts {*x*_*t*_} in 10 ms bins at the trajectory positions. We then ran our inference pipeline on each SWR, recovered estimates 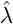 and 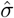 and compared them to the ground truth.

##### Switching of dynamics (dynamical regimes)

To test recovery of both regime structure and drift-diffusion parameters, we expanded our ground truth to: (a) every ordered pair of candidate dynamics (drift-diffusion, stationary, uniform), and (b) for any segment governed by drift-diffusion, each (*λ*_gt_, *σ*_gt_) combination from the same grid as in drift-diffusion regime experiment. For each of these regime-pair *×* parameter conditions, we simulated 100 independent SWRs as follows. First, specify the two-segment regime sequence (e.g., drift-diffusion → uniform). Next, generated a latent path for each 300 ms segment. Use the drift-diffusion dynamics with the given *λ*_gt_ and *σ*_gt_ when applicable, or the stationary/uniform dynamics otherwise. Concatenate the two segments into a continuous 600 ms trajectory. Along this trajectory, sample Poisson spike counts from place-cell population in 10 ms bins. By applying the inference pipeline to each simulated SWR, we recovered the latent regime sequence {*ĉ*_*t*_} and, when drift–diffusion segment presents, the parameter estimates 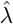 and 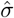. We then compared {*ĉ*_*t*_}, 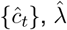 and 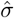 against the ground-truth.

#### Comparative analysis between drift-diffusion parameters and prior replay metrics

We simulated SWR events under the pure drift-diffusion dynamics using the same settings as in identifiability experiments. We ran two sets of simulations: (a) Fix the 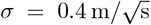 and simulate 2000 SWRs for each *λ*_gt_ ∈ {0, 2, 4, 6, 8, 10} m/s. (b) Fixed the *λ* = 2 m/s and simulate 2000 SWRs for each 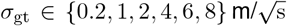. We applied classic Bayesian decoding to calculate the location posterior and MAP trajectory. From these decoded trajectories, we then computed absolute correlation, maximum jump distance and replay speed as outlined above, and compared them to the ground-truth *λ*_gt_ and *σ*_gt_.

### 4.7 Additional details of the figures

#### Speed calculation

We computed three complementary measures of speed in Fig. 6. *Per-lap running speed*. Individual laps on the linear track were identified by crossings of the track midpoint (90 cm). For an up-crossing of the midpoint, the start of the lap was defined as the most recent up-crossing of 65 cm, and the end as the first new up-crossing of 115 cm. For down-crossings, the similar down-crossings of 115 cm and 65 cm were used. After the position-time trace of each lap was extracted, we fit a simple linear regression to the data within that lap and took the per-lap running speed as the absolute value of slope. *Instantaneous behavioral speed*. Within each running epoch, we estimated instantaneous velocity by two-step centered differencing of the position time series, *v*(*t*) = (pos(*t* + 1) − pos(*t* − 1)) */*2Δ*t*, where Δ*t* = 10 ms is the time bin. The instantaneous behavioral speed was taken as the absolute value |*v*(*t*)|. *Replay speed*. We used the absolute value of inferred drift-rate parameter |*λ*| (in m/s) as replay speed.

#### Shuffle analysis

For Fig. 8, null replay distributions were generated using three types of shuffle procedures. *Time-bin shuffle*. The time bins of spike raster were randomly permuted within each SWR. *Neuron-identity shuffle*. The neuron indices of spike raster were randomly permuted within each SWR. *Place-field shuffle*. Place fields of each neuron were circularly shifted by a random offset. Each shuffle procedure was applied to every SWR and repeated 120 times to construct a null distribution. Shuffled SWRs were then processed and analyzed using the same pipeline used for real replay events.

### 4.8 Data and code availability

This study uses data from Pfeiffer and Foster (2015) [33]. Please contact Brad Pfeiffer and David Foster to access the analyzed datasets. The code used for data analysis, simulations, and figure generation is archived on a GitHub repository (https://github.com/ZhongxuanWu/ReplaySwitchingHMM). Any further information needed to reproduce the results can be obtained from the lead contact upon request.

## Supplemental Information

## A Additional figures and discussions

### A.1 Parameters in drift-diffusion framework are more interpretable than prior metrics

Previous methods first infer the posterior probability over space using the place fields inferred during running [1], then compute three metrics for interpretation (summarized in [2]): (i) (absolute) correlation between time and decoded trajectory; (ii) replay speed inferred from the slope of decoded trajectory; and (iii) maximum jump distance (Fig. S1A). In contrast, our modeling framework uses a generative approach and describes a replay event using the drift and diffusion parameters, which unambiguously characterize the speed and the sequentiality (or quality) of the replay respectively. We next aim to determine the relationship between the key parameters in prior methods and those in ours.

We generate simulated neural population recording data with different combinations of drift and diffusion parameters, and then apply the traditional analysis method based on Bayesian decoding to extract the three metrics mentioned above, i.e., absolute correlation, speed of replay, maximum jump distance (Fig. S1A). Fig. S1B shows the results when fixing the diffusion parameter while allowing the drift parameter to vary, while Fig. S1C shows the results when fixing the drift parameter while allowing the diffusion parameter to vary. These results suggest that the absolute correlation is positively correlated with drift and negatively correlated with diffusion. For the speed inferred from the Bayesian approach, it is positively correlated with both the drift and diffusion parameter. Thus, the speed can be overestimated when the diffusion parameter is large. Furthermore, the maximum jump distance [2] mostly reflects the magnitude of the diffusion parameter, i.e., large diffusion leads to bigger maximum jump distance, yet it also increases systematically with the drift. This suggests that maximum jump distance reflects a combination of drift and diffusion, in particular when the diffusion parameter is small. Together, these results suggest that, while the three key metrics used in prior studies may capture some structures in the replay dynamics as conceived when they were developed, each of them reflects the combined effects of the speed and the quality of a candidate replay event. Thus, practically they may not allow for an unambiguous interpretation of the replay dynamics. In contrast, our approach clearly distinguishes the speed and quality of a replay event using separate drift and diffusion parameters, and it allows one to dissect the contribution of the two in a principled single-step inference framework.

### A.2 Discussion of different shuffle controls when analyzing pre-running SWRs

Across shuffle controls when analyzing pre-running SWRs, we observed a specific discrepancy for the time-bin shuffle (Fig. 8 and Fig. S8). First, the fraction of SWRs exceeding each drift-diffusion duration threshold for time-bin shuffled data was consistently lower than that for the other two controls, whereas the neuron-identity and place-field rotation shuffles yielded similar null fractions. Second, the null fractions from time-bin shuffle did not align across pre- and post-run data. At every threshold a larger fraction of post-run shuffled events exceeded the criterion than pre-run shuffled events did. This is in contrast to the overlapping pre/post null fractions produced by the neuron-identity and place-field rotation shuffles.

Note that a comparable divergence between time-bin and other shuffles was also reported by Takigawa et al. [3]. The reason is possibly that only the time-bin shuffle preserves spike–position correlations [4, 5]. But this interpretation is provisional, and the mechanism requires further investigation.

### A.3 Discussion of bidirectional place fields

On a linear track, place cells may exhibit direction-dependent place fields. In our analyses, we estimate place fields of each neuron by pooling spikes and occupancy from both running directions. For clarity, we refer to these pooled estimates as bidirectional place fields throughout and use them in the main analyses. We chose to combine the data from the two directions for the following reasons. First, the peaks of individual place fields are highly consistent across directions (Fig. S9 and Fig. S11). Second, using the bidirectional fields, the animal’s position during run periods can be decoded accurately (Fig. S10 and Fig. S12). Therefore, if replay recapitulates place cell activity during running and forms coherent trajectories, the bidirectional place fields should be appropriate for decoding the content of the replay events. Finally, our choice matches the preprocessing in the original report on this dataset [6], which did not split place fields by running directions.

**Figure S1:**
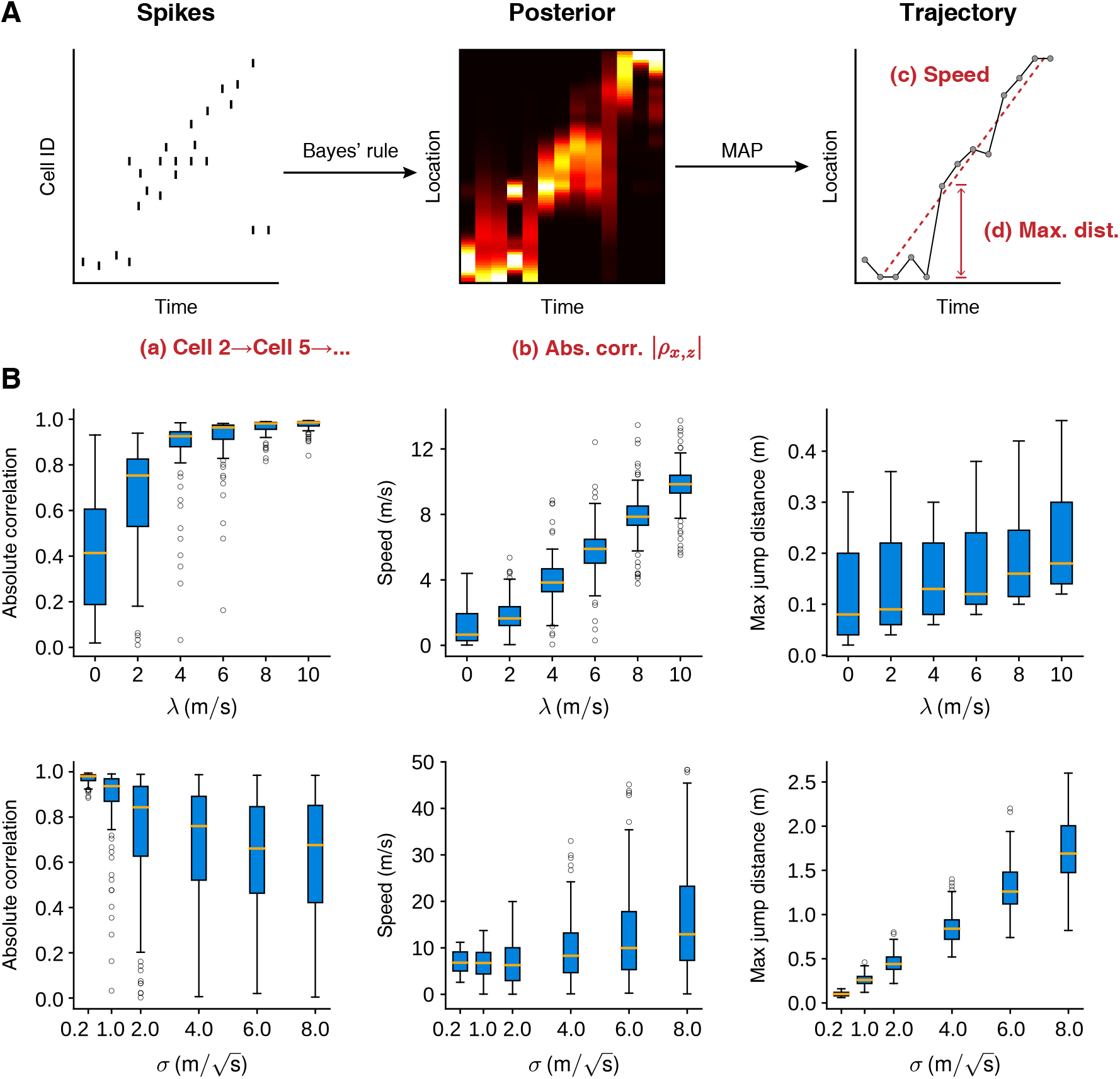
Relationship to previous replay analysis framework. **A.** Previous two-step replay analysis. Bayesian decoding (in black). Left: Spike-raster matrix for a single SWR. Middle: Bayesian location posterior *p*(*z*_*t*_|*x*_*t*_) ∝ *p*(*x*_*t*_|*z*_*t*_)*p*(*z*_*t*_), computed using each neuron’s place field and a uniform prior *p*(*z*). Right: decoded trajectory obtained as the MAP estimate 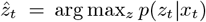. Metric calculation (in red). Temporal firing sequence, compared against the animal’s running firing template. Absolute correlation |*ρ*_*t,z*_ |: weighted Pearson correlation between time bin *t* and position *z* under the posterior, reflecting sequence linearity. Replay speed: absolute slope |*β*_1_| of a linear regression fit 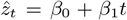. Max jump distance: 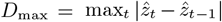, the largest spatial jump between consecutive bins. **B**. Boxplots illustrate how the three classic Bayesian replay metrics depend on the drift *λ* (top row) and diffusion parameter *σ* (bottom row). In each plot, one parameter is held constant while the other varies. For *λ* sweeps, 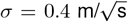 and for *σ* sweeps, *λ* = 2 m/s. Boxes show the median and interquartile range, and whiskers extend to 1.5*×* IQR.

**Figure S2:**
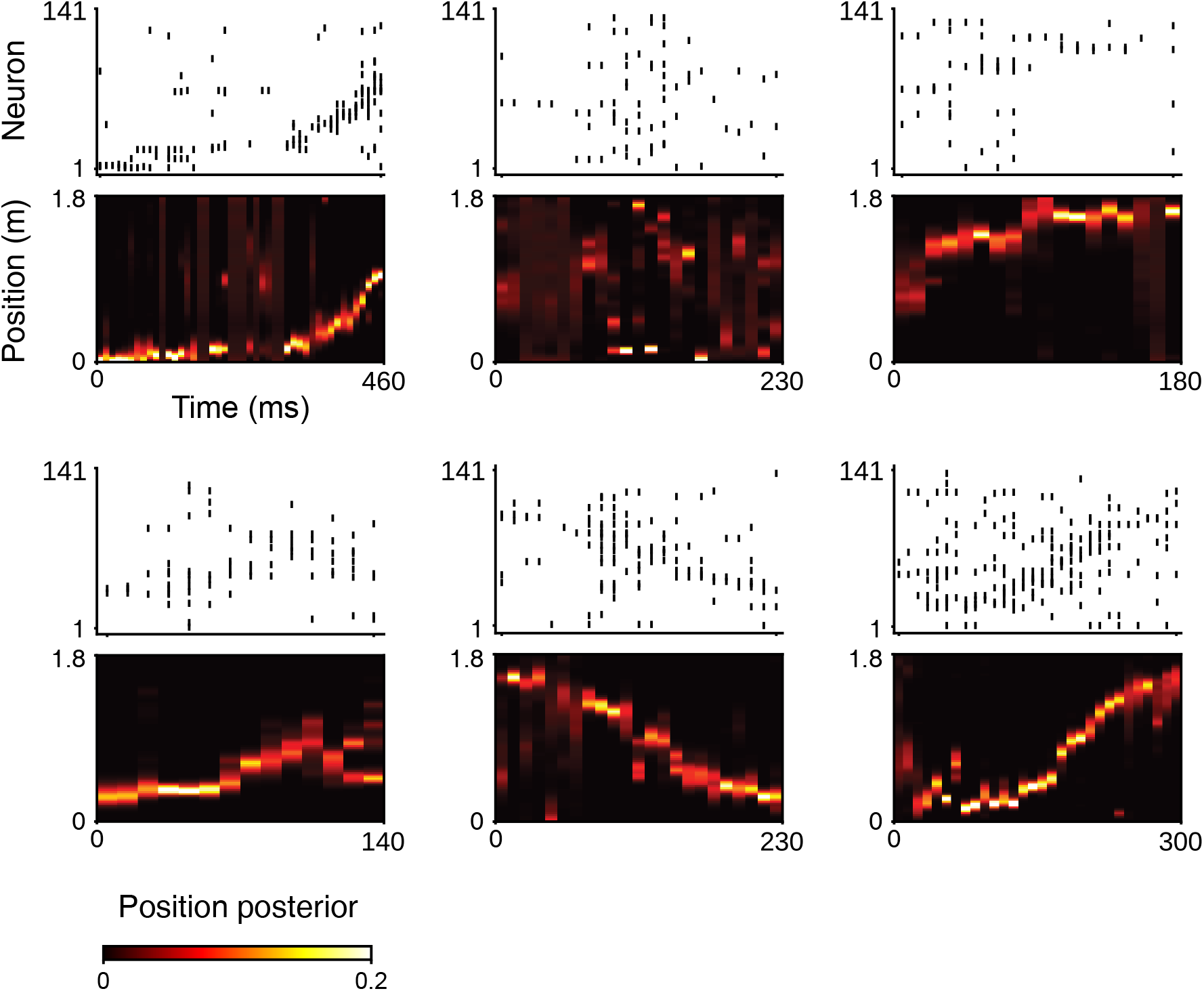
Spike rasters and position posteriors from classic Bayesian decoding for the example events. These events correspond to those shown in Fig. 6A. For each event, the spike raster (top, neurons sorted by place field peak location) and decoded position posterior (bottom) are shown. Posteriors were computed with a Poisson Bayesian decoder using 10 ms time bins and no temporal smoothing. Horizontal axis: time (ms). Vertical axis: linear-track position (m). Color bar: posterior probability.

**Figure S3:**
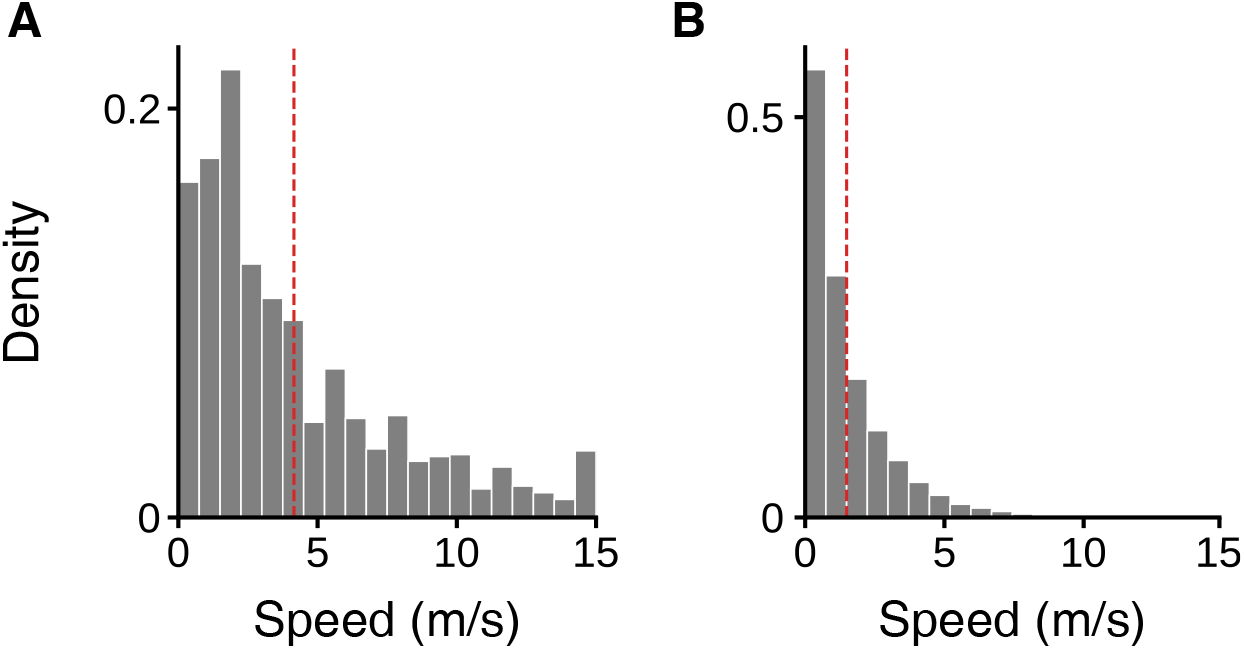
Replay-speed distribution observed in data is inconsistent with a pure Brownian motion null. **A.** Histogram of replay speed, estimated by the inferred drift rate *λ* for SWRs in which the marginal posterior of drift-diffusion regime exceeded 0.7 for at least 50 ms. Panel repeats the one shown in Fig. 6 for comparison. **B**. Histogram under a pure Brownian motion null. For each SWR in A, we fit a Brownian motion model by MLE to estimate the diffusion parameter and simulate 10 trajectories of matching duration. For each trajectory, we compute its speed as the absolute value of slope of a linear regression fit to position and time. The pooled distribution of speeds is shown. Dashed vertical lines show the mean of each distribution: 4.1 m/s (replay) and 1.5 m/s (null).

**Figure S4:**
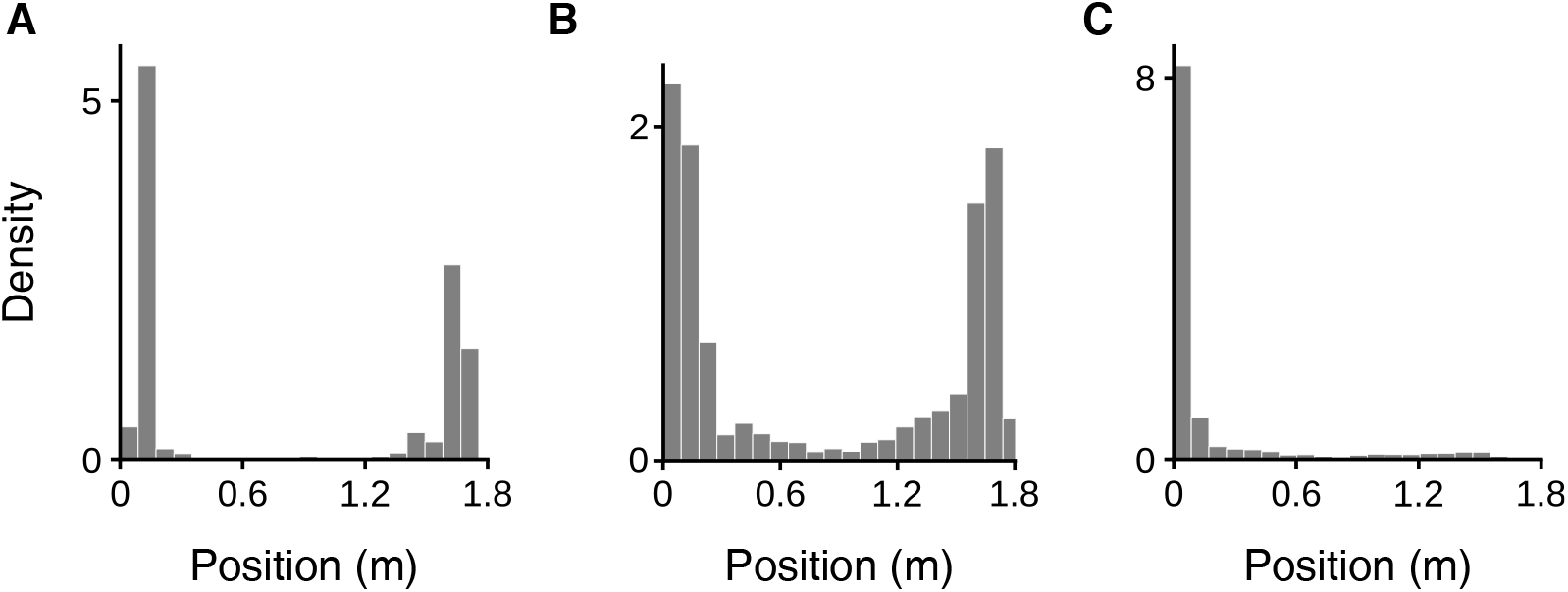
The distribution of latent position during stationary periods. Analyses combined periods that our model labeled as stationary within awake-rest SWR events (total duration *T* = 88.99 sec across 9 sessions). **A**. Histogram of the actual position of rats on the linear track during awake SWR periods. **B**. Histogram of the decoded latent position. **C**. Histogram of the absolute difference between actual and decoded latent positions.

**Figure S5:**
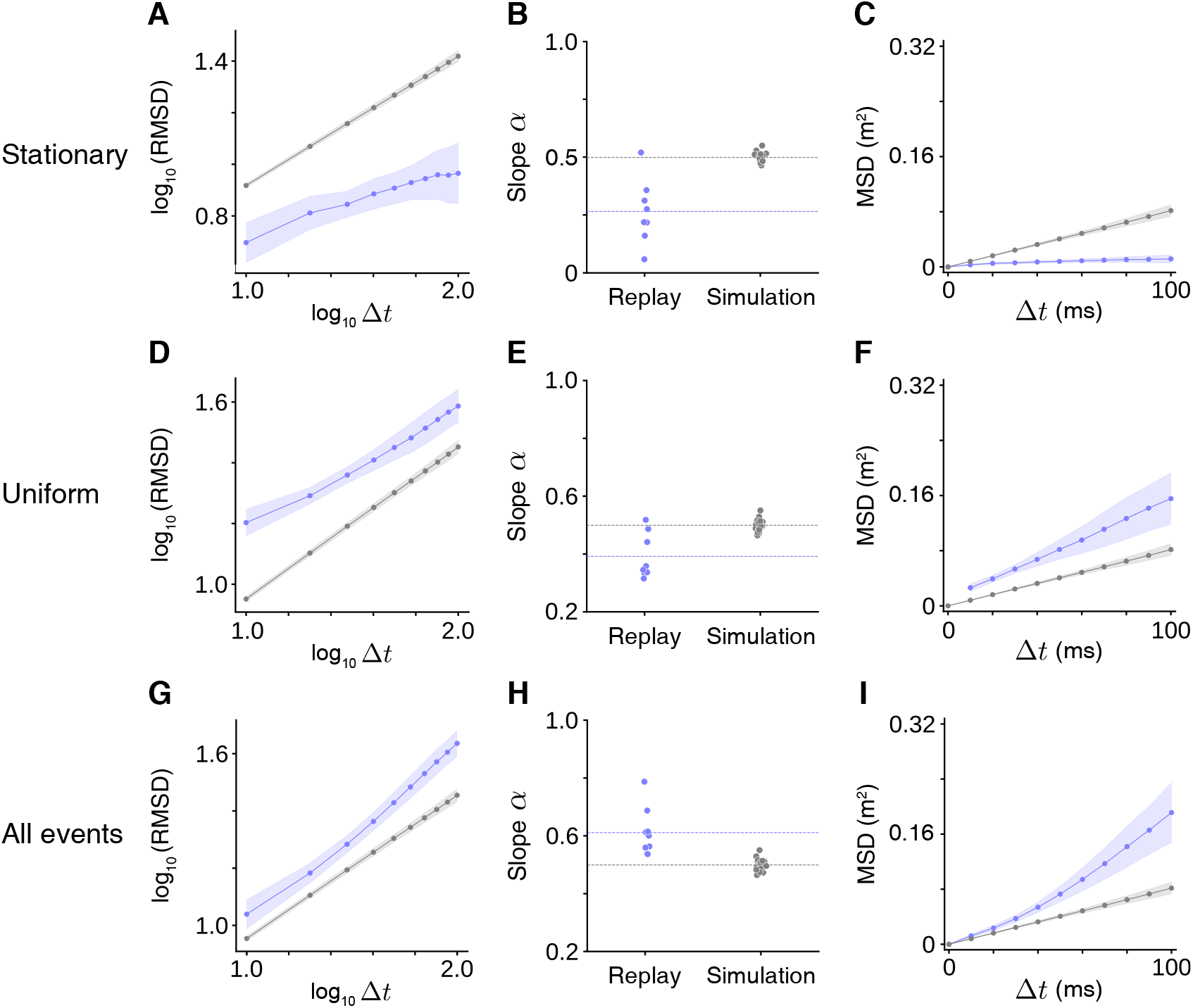
The scaling relationships of RMSD-Δ*t* and MSD-Δ*t* are distinct for stationary, uniform, and all SWR events. A-C. Plotting conventions follow Fig. 7B-D. Analyses repeated to SWR events with pure stationary dynamics. The mean across sessions in **B**: 0.26 for SWRs with stationary dynamics; 0.50 for simulated data from Brownian motion. **D-F**. Analyses repeated to SWR events with pure uniform dynamics. The mean across sessions in **E**: 0.39 for replay; 0.50 for simulation. For **F**, the real-data point at Δ*t* = 0 is omitted. The reason is that for uniform dynamics, the MSD is strictly positive for any Δ*t* ≥ 0 but the value at Δ*t* = 0 is degenerate and cannot be estimated reliably. **G-I**. Analyses repeated to all SWR events. The mean across sessions in **H**: 0.61 for replay; 0.50 for simulation. One session was excluded from the analyses in **A-F** because fewer than 50 SWR events met the inclusion criteria.

**Figure S6:**
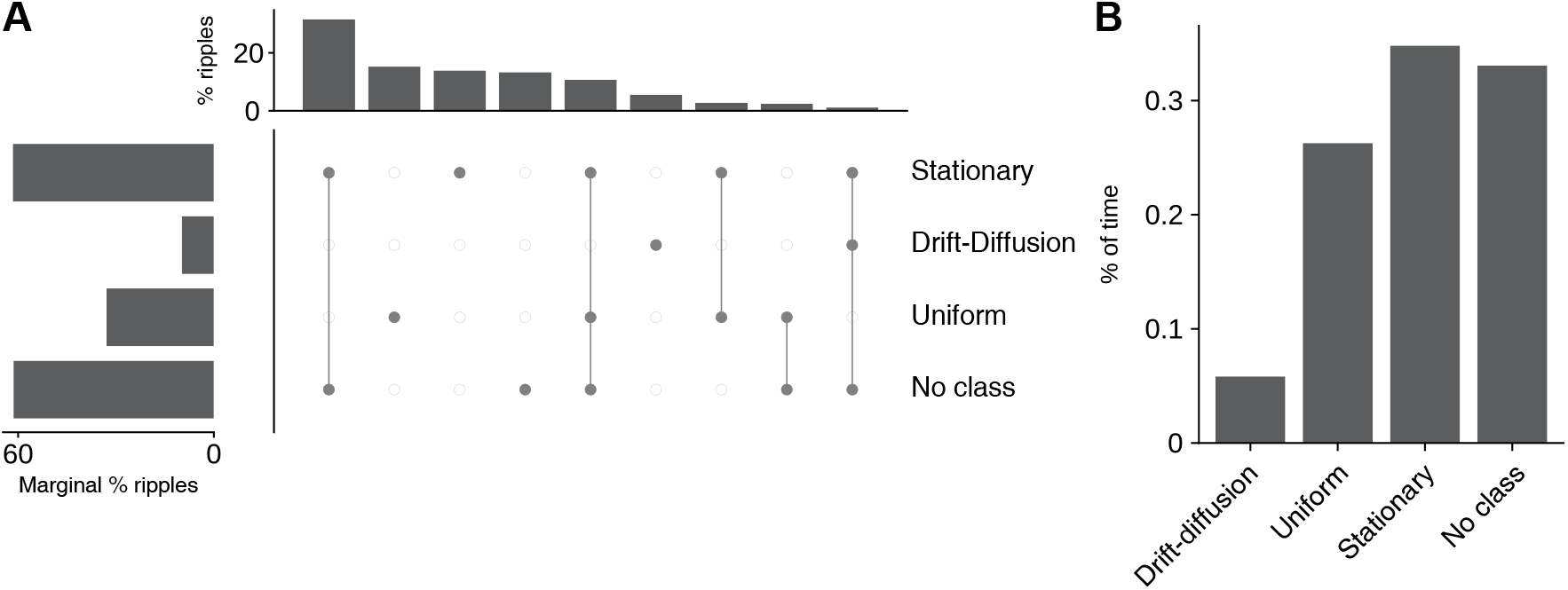
Composition of spatiotemporal dynamics of pre-run SWRs. A-B. Plotting conventions follow Fig. 4. Compared to the post-run, pre-run SWRs showed a markedly lower occurrence of drift-diffusion dynamics and a higher occurrence of uniform dynamics.

**Figure S7:**
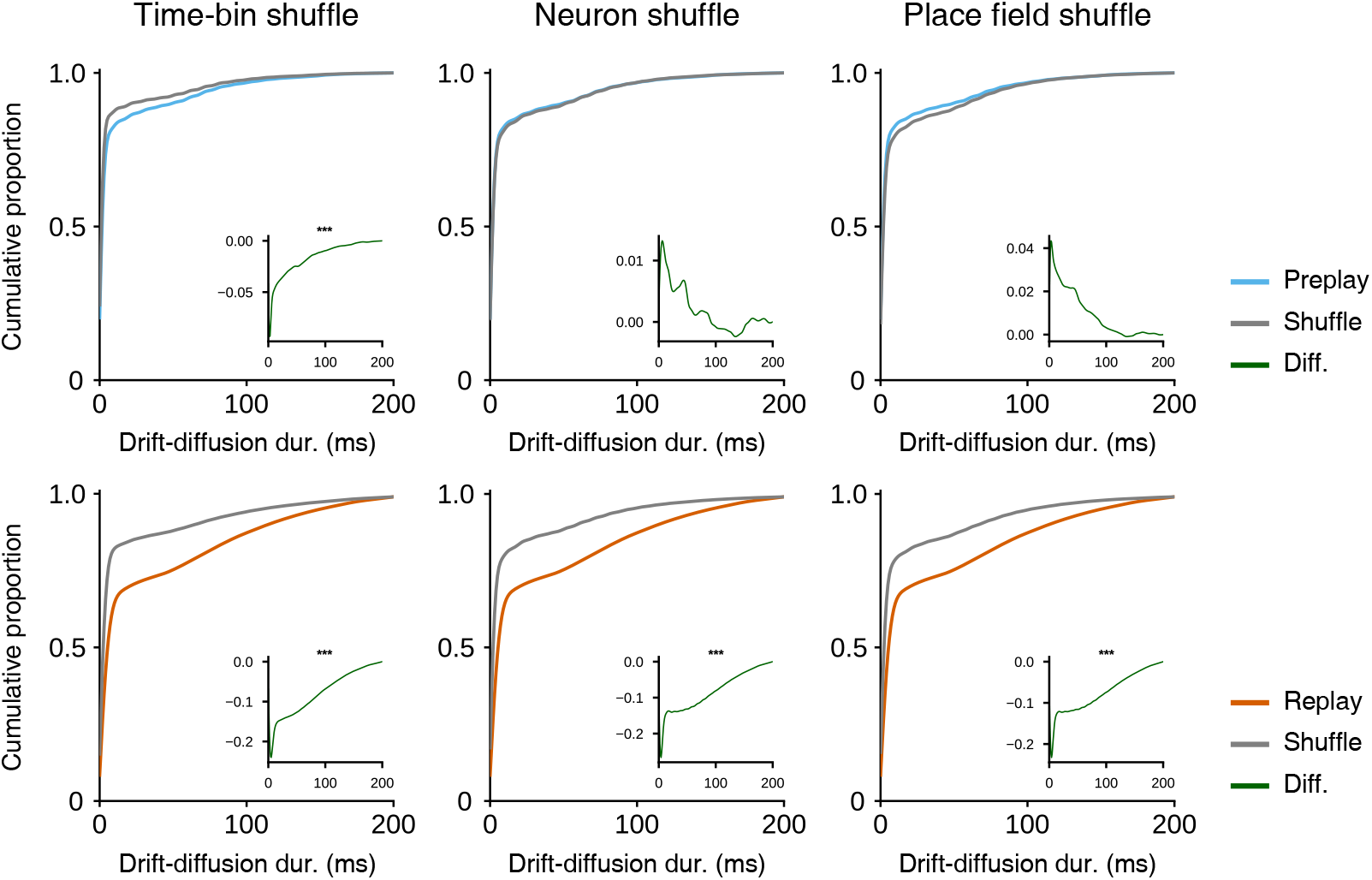
Comparison of drift-diffusion duration in preplay and replay SWRs to corresponding shuffle. Cumulative proportions of drift-diffusion durations for pre-run (top row, blue) and post-run (bottom row, orange) SWRs, compared to shuffled SWRs. Columns correspond to three shuffle controls—time-bin (left), neuron-identity (middle), and place-field rotation (right) Inset: the point-wise difference between the two cumulative proportions. Asterisks mark a significant divergence between distributions (one-sided Kolmogorov–Smirnov test, *** *p <* 0.001).

**Figure S8:**
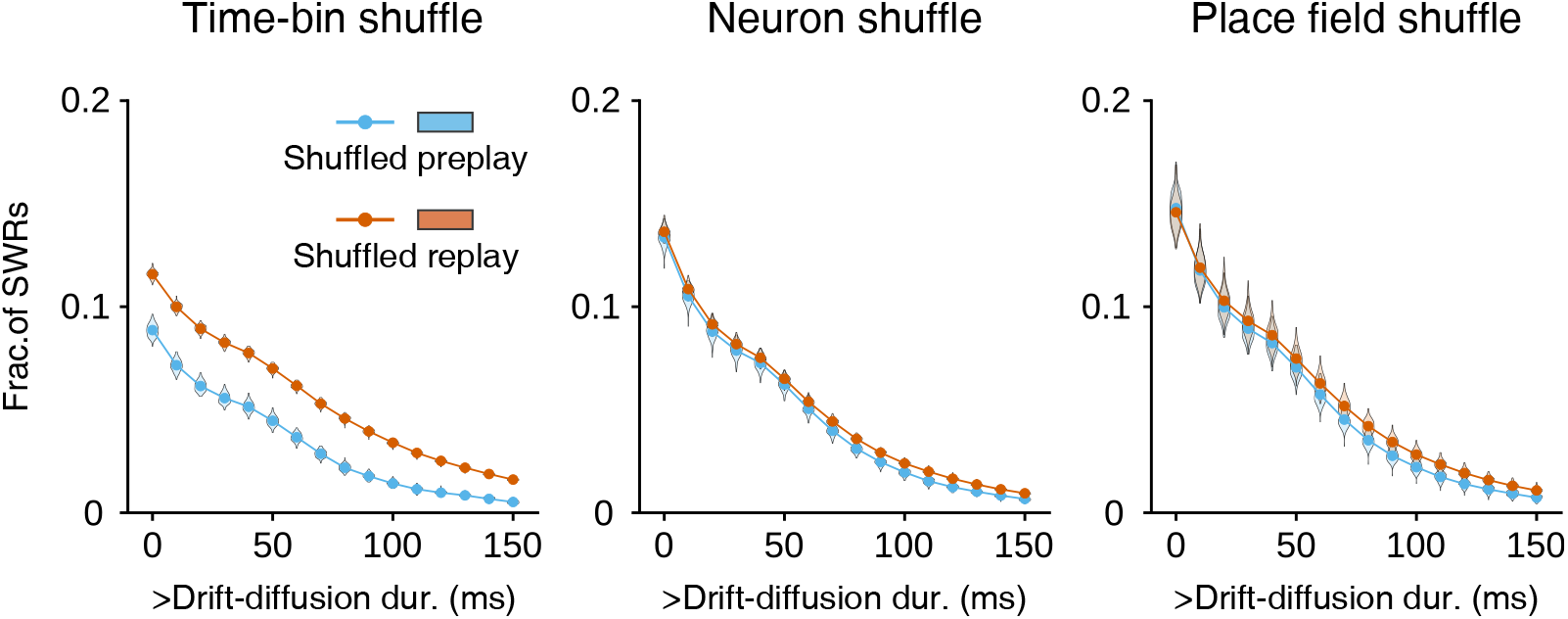
Comparison of shuffled preplay versus shuffled replay SWRs. The fraction of shuffled preplay (blue) and shuffled replay (orange) SWRs exceeding each drift-diffusion duration threshold (calculation as in Fig. 8A-F). Solid lines plot the mean fractions. Violins plot the distribution of the fractions across 120 shuffles. Columns show three shuffle methods: time-bin (left), neuron-identity (middle), and place-field rotation (right).

**Figure S9:**
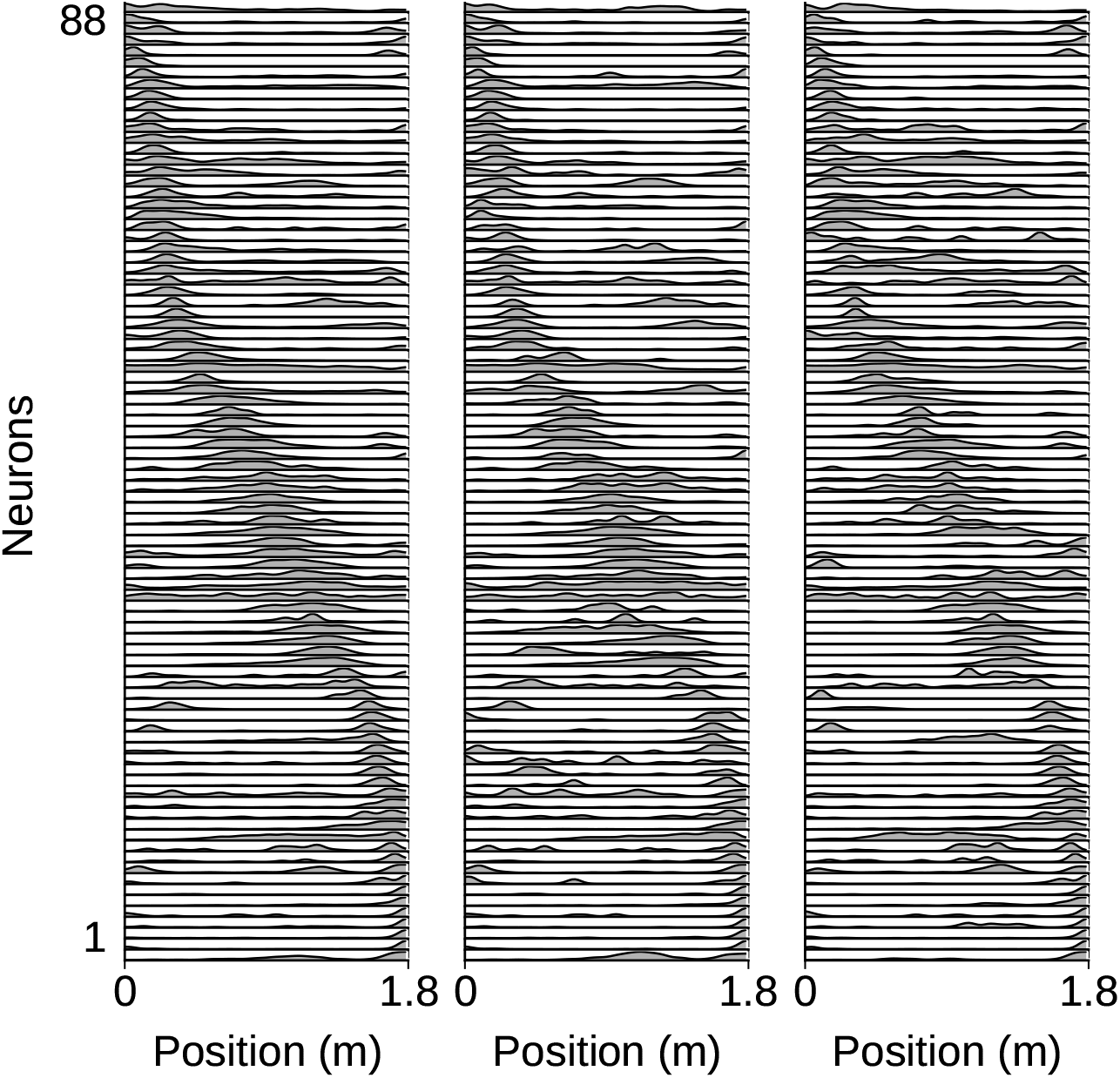
Place fields in an example session of rat Janni. Left: bidirectional place fields obtained by pooling spikes and occupancy from both running directions. Middle: place fields estimated from left-to-right traversals only. Right: place fields estimated from right-to-left traversals only. Each row is the place fields of one neuron. Place fields are normalized to the peak firing rate. Neurons are ordered by the peak position of the bidirectional place fields, and the same order is used in all panels.

**Figure S10:**
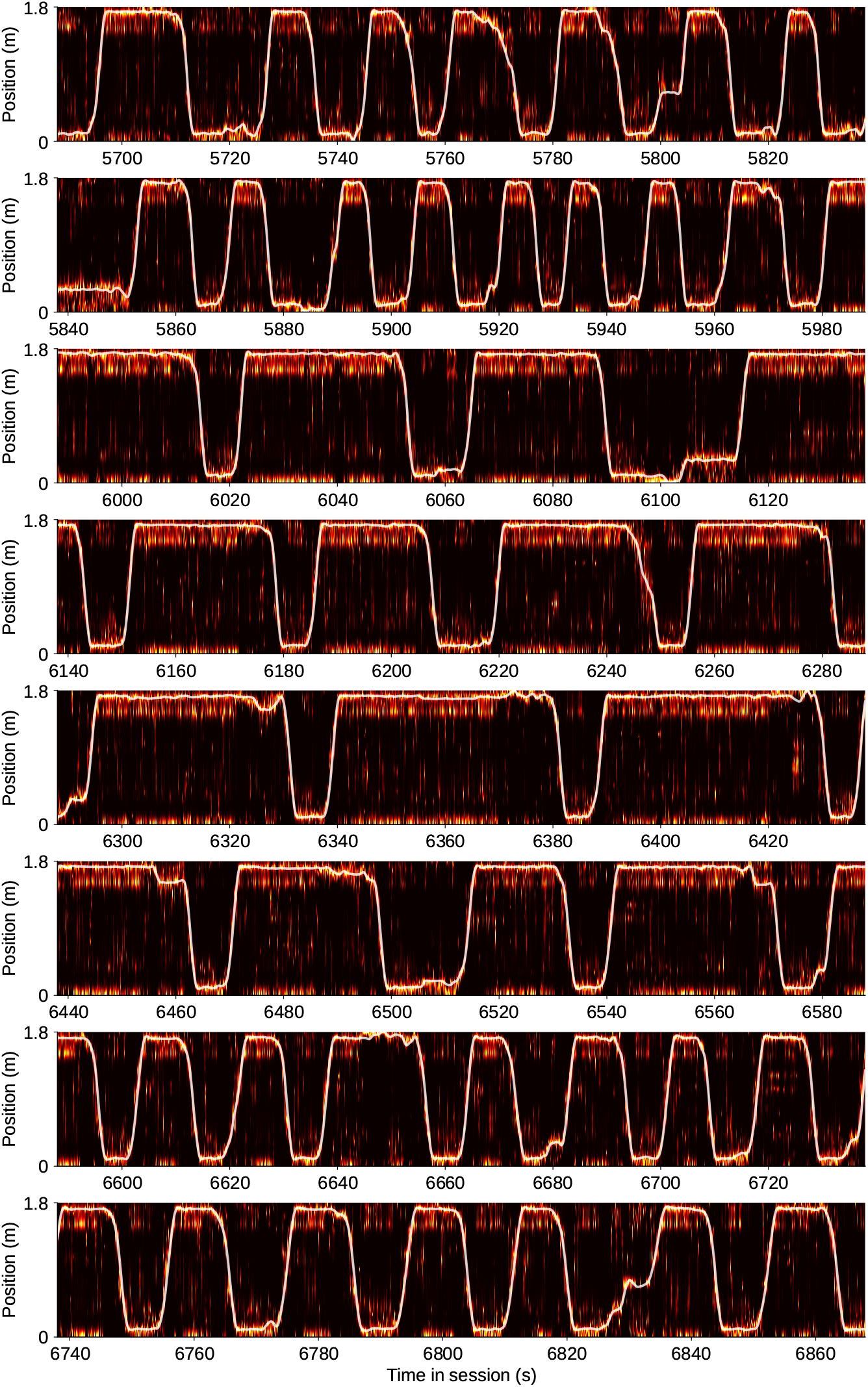
Decoding run position using bidirectional place fields. Heatmap shows the position posterior along the linear track (brighter = more probable). This is estimated by a memoryless Bayesian decoder [1] applied to place cell activity during the run epoch. The white trace shows the tracked position of the animal. Panels depict consecutive time segments from the same session. Same session as in Fig. S9.

**Figure S11:**
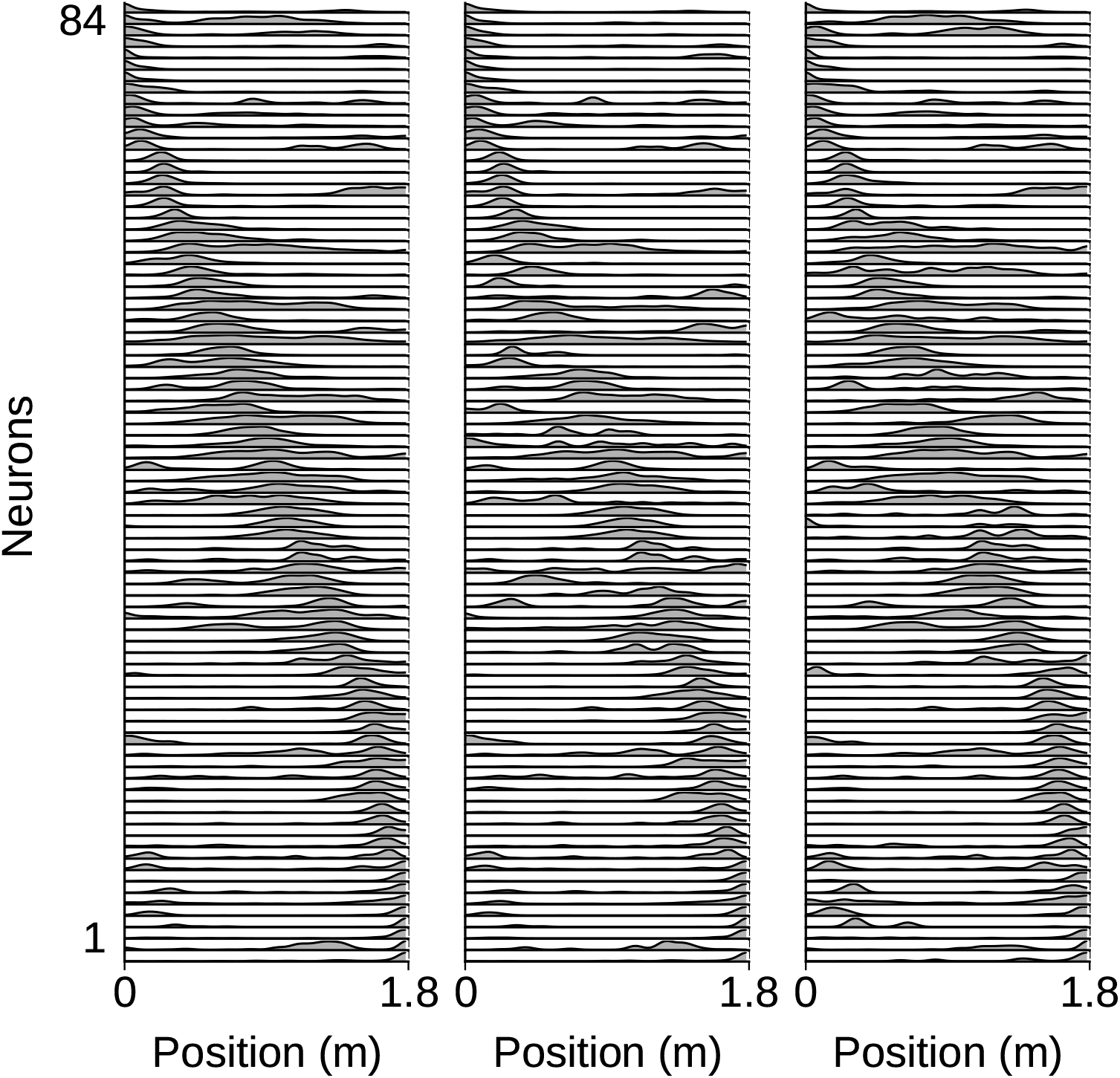
Place fields in an example session of rat Harpy. Plotting conventions follow Fig. S9.

**Figure S12:**
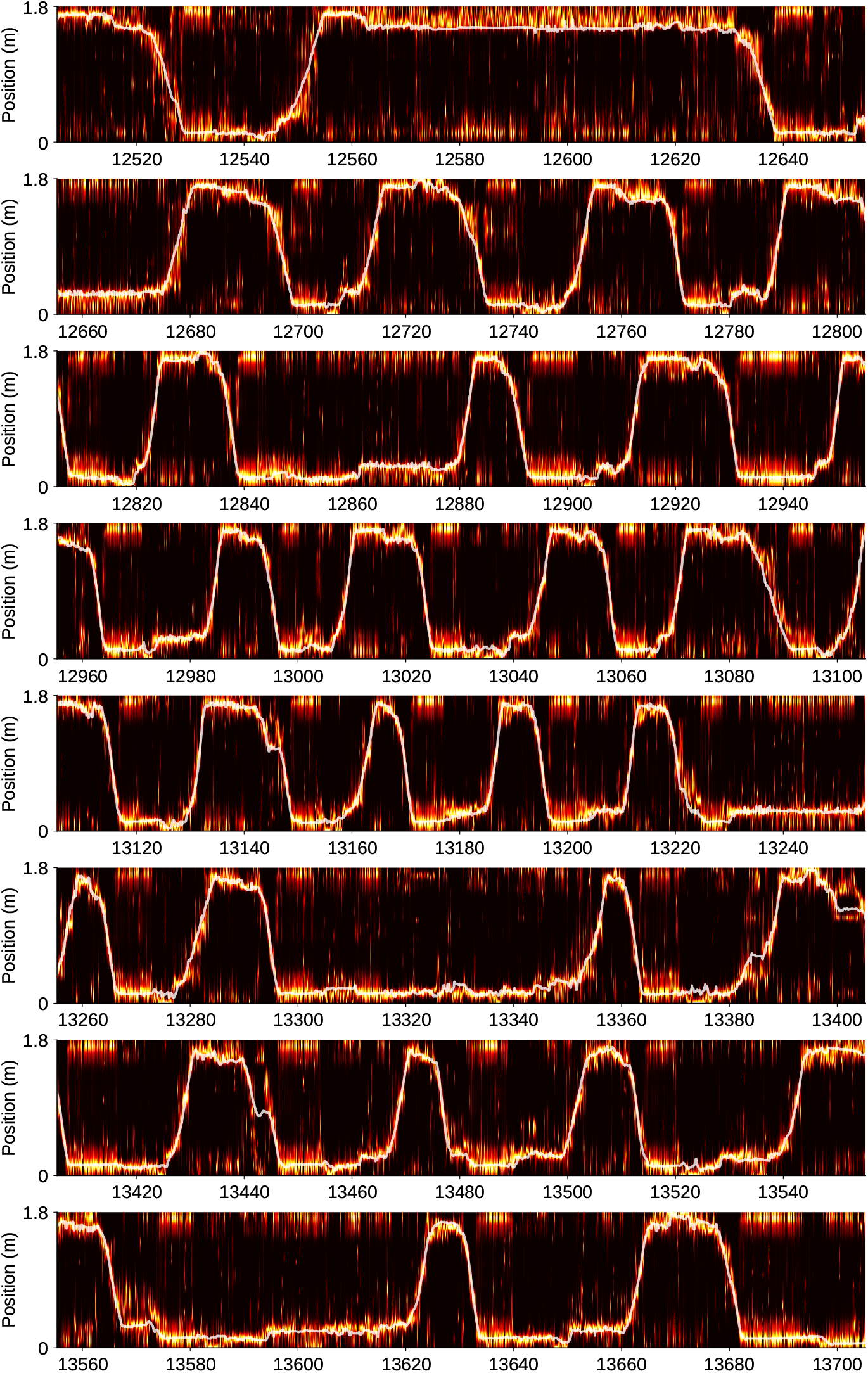
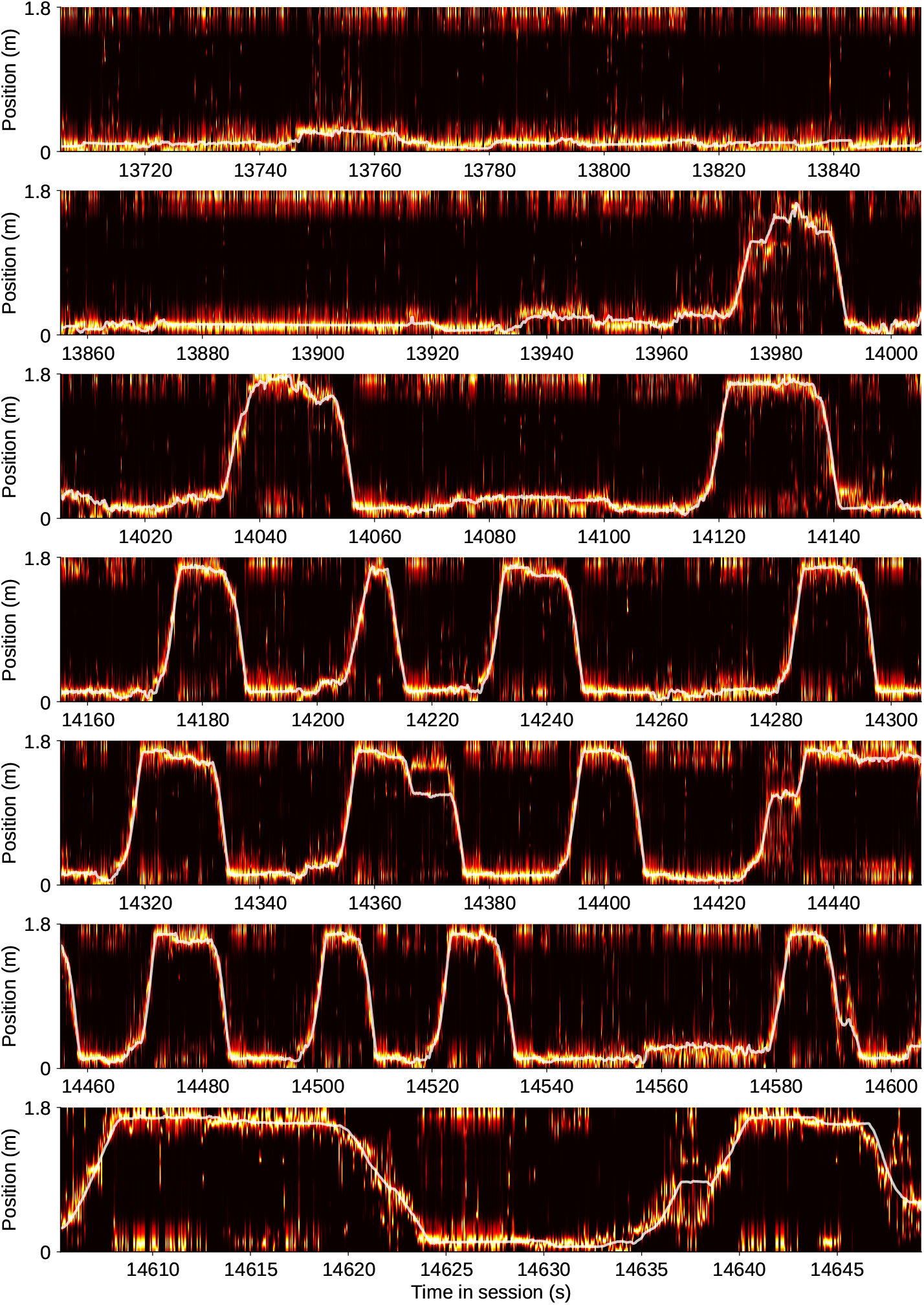
Decoding run position using bidirectional place fields. Plotting conventions follow Fig. S10. Same session as in Fig. S11.

## B Mathematical notes on the model

### B.1 Derivation of posterior of the switching HMMs

We consider a regime-switching hidden Markov model. At each discrete time point *t*, an observed variable *x*_*t*_ ∈ 𝒳 is generated by a latent Markov chain *z*_*t*_ ∈ {1, …, *S*}, whose transition probabilities are modulated by a discrete regime variable *c*_*t*_ ∈ {1, …, *C*}. The joint evolution is specified by

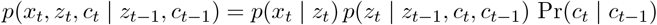

where *p*(*x*_*t*_ | *z*_*t*_) is the emission distribution, *p*(*z*_*t*_ | *z*_*t*−1_, *c*_*t*_, *c*_*t*−1_) is the state-transition kernel conditional on current and previous regime, and Pr(*c*_*t*_ | *c*_*t*−1_) is the regime-switching matrix.

#### Forward pass

We consider the forward message,

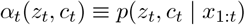

By Bayes’ rule,

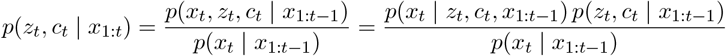

The Markovian assumption and the emission independence give *p*(*x*_*t*_ | *z*_*t*_, *c*_*t*_, *x*_1:*t*−1_) = *p*(*x*_*t*_ | *z*_*t*_, *c*_*t*_) = *p*(*x*_*t*_ | *z*_*t*_). Thus,

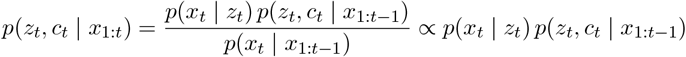

Using the Chapman-Kolmogorov equation, we can express the one-step prior *p*(*z*_*t*_, *c*_*t*_ | *x*_1:*t*−1_) in terms of *α*_*t*−1_. By marginalizing over the previous latent pair (*z*_*t*−1_, *c*_*t*−1_),

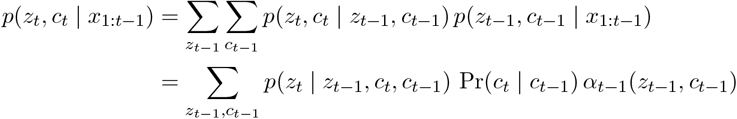

Combining the prior into the Bayes proportionality gives the usual recursion:

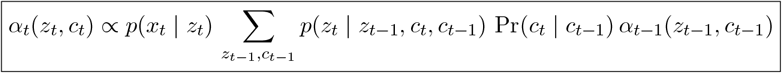

The initialization of *α*_*t*_(*z*_*t*_, *c*_*t*_) can be specified as *α*_1_(*z*_1_, *c*_1_) ∝ *p*(*x*_1_ | *z*_1_) *π*_*z*_(*z*_1_ | *c*_1_) *π*_*c*_(*c*_1_), where *π*_*c*_ and *π*_*z*_(· | *c*_1_) are the regime and state priors. The normalizing constant is 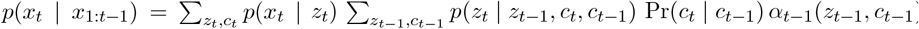 so that 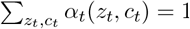.

#### Backward pass

We consider the backward message,

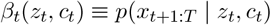

By marginalizing over the next latent pair (*z*_*t*+1_, *c*_*t*+1_),

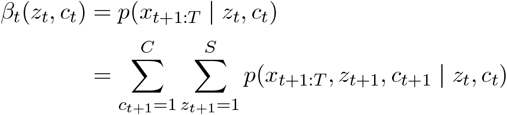

Apply the chain rule to factorize,

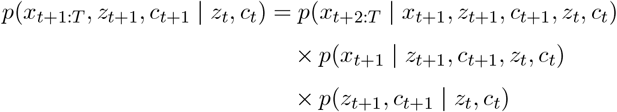

The future observations *x*_*t*+2:*T*_ are conditionally independent of (*z*_*t*_, *c*_*t*_, *x*_*t*+1_) given (*z*_*t*+1_, *c*_*t*+1_),

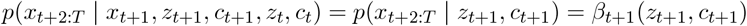

The emission depends only on *z*_*t*+1_, not on *c*_*t*+1_ or past states,

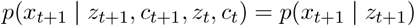

The joint transition factorizes as

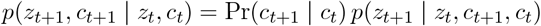

Combine and obtain the recursion,

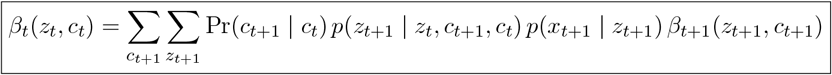

The initialization is *β*_*T*_ (*z*_*T*_, *c*_*T*_) = 1 for all (*z*_*T*_, *c*_*T*_).

#### Model posterior

At time *t*, the posterior is defined as

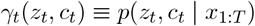

Express it as the full joint,

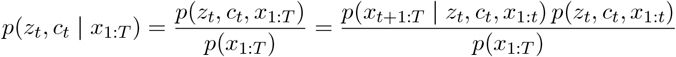

By the Markov structure,

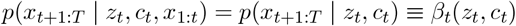

Likewise,

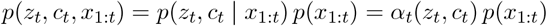

Putting these together,

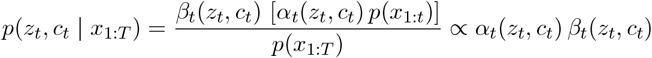

Or

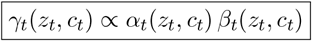

where the constant of proportionality *p*(*x*_1:*t*_)*/p*(*x*_1:*T*_) is fixed by 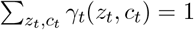.

### B.2 Discussion on the estimation bias of drift and diffusion parameters

For simplicity, we perform analysis based on HMM, and briefly discuss switching HMM later. Let the 1-D track be the closed interval with grid points *z*_*t*_ ∈ {1, …, *S*} and spacing Δ*z*. Time is discretized in steps of size Δ*t*. The transition matrix is parameterized by a drift-diffusion step:

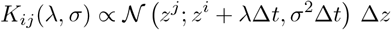

where 𝒩 (·; *µ, ν*) is the Gaussian density, *λ* is the drift parameter, and *σ* is the diffusion scale. Observations **x**_0:*T*_ are replay spike trains. The emission probabilities *p*(**x**_*t*_ | *z*_*t*_ = *z*^*j*^) = *B*_*j*_(**x**_*t*_) are specified by place fields fit from running data. The marginal likelihood *L*(*λ, σ*) is evaluated with the forward algorithm, and we maximize log-likelihood *ℓ*(*λ, σ*) = log *L*(*λ, σ*) to obtain the estimated parameters. For notation in derivatives, we use the usual marginal probabilities obtained via forward-backward algorithm:

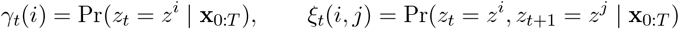

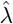 **is unbiased**. Let *ℓ*(*θ*) = log *p*(**x**_0:*T*_ | *θ*) with *θ* = (*λ, σ*). We start from,

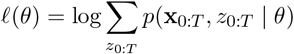

Differentiate using ∂_*λ*_*p* = *p* ∂_*λ*_ log *p*:

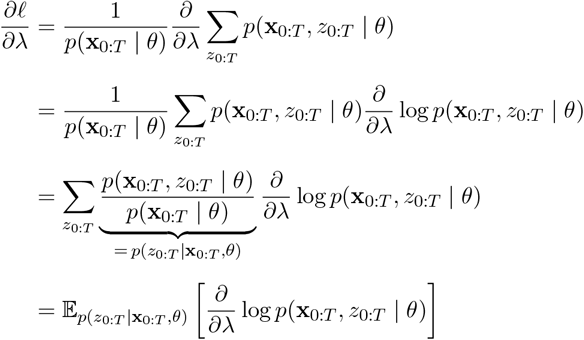

The complete-data joint distribution factors as,

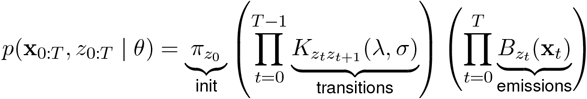

Taking logs,

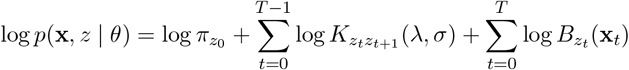

When differentiating w.r.t. *λ*,

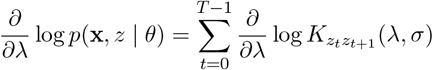

Plug this into the expectation:

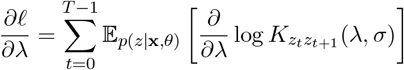

Write the expectation using the pairwise marginals *ξ*_*t*_(*i, j*) = Pr(*z*_*t*_ = *i, z*_*t*+1_ = *j* | **x**_0:*T*_, *θ*) obtained by forward–backward:

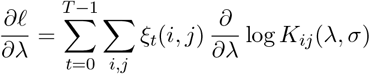

For a Gaussian transition matrix,

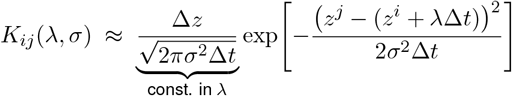

Hence,

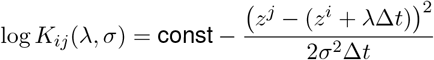

Differentiate w.r.t. *λ*. Let *µ*_*i*_(*λ*) = *z*^*i*^ + *λ*Δ*t* and *v* = *σ*^2^Δ*t*.

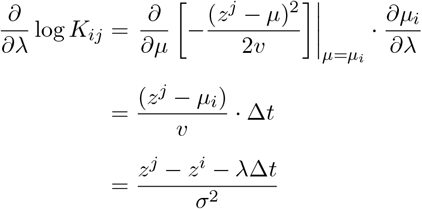

Therefore,

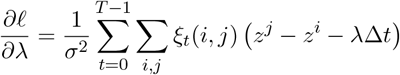

Setting the score function to 0 gives,

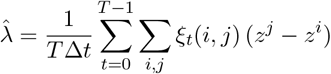

Using *ξ*_*t*_(*i, j*) = Pr(*z*_*t*_ = *z*^*i*^, *z*_*t*+1_ = *z*^*j*^ | **x**_0:*T*_, *θ*),

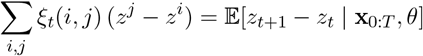

If we evaluate the posteriors at the true parameters (*λ*^∗^, *σ*^∗^), by the law of iterated expectations,

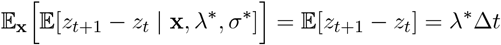

Hence,

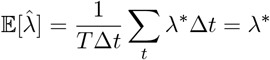

This means the MLE estimate has zero bias.

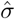 **is biased downward**. With *θ* = (*λ, σ*) and *ℓ*(*θ*) = log *p*(**x**_0:*T*_ | *θ*),

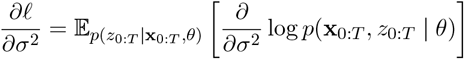

Using the factorization,

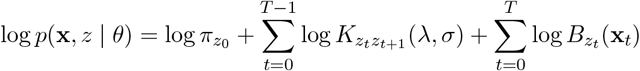

Only the transition terms depend on *σ*. Hence,

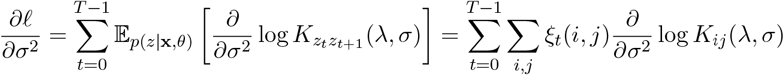

With Gaussian transition kernel,

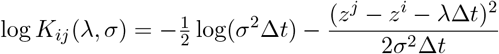

Let *A*_*ij*_ := *z*^*j*^ − *z*^*i*^ − *λ*Δ*t*. Then differentiate log *K*_*ij*_ w.r.t. *σ*^2^,

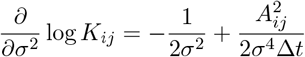

Set the score to zero and solve,

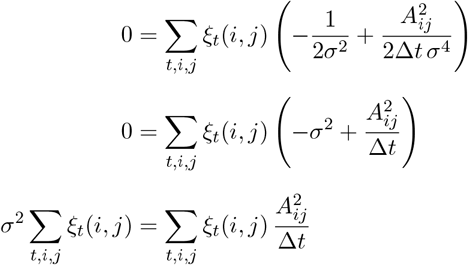

Hence we have the variance MLE estimator,

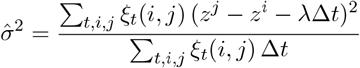

Using ∑_*i,j*_ *ξ*_*t*_(*i, j*) = 1,

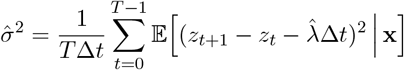

If *λ* were known and 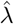 is equal to *λ*^∗^, then by the law of iterated expectations,

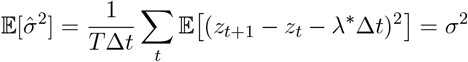

If 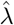 is estimated from the same data, we evaluate at the true parameters (*λ*^∗^, (*σ*^∗^)^2^).

Define increments

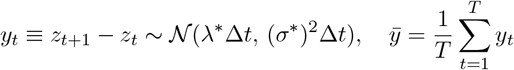

For simplicity, we further approximate 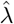 as 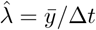 under the assumption that emissions are informative and *ξ*_*t*_(*i, j*) are sharply peaked. We then calculate moments,

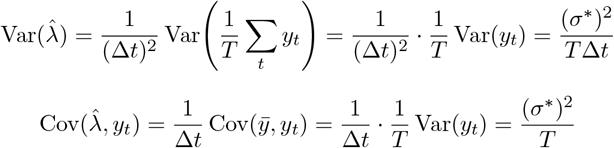

By the law of iterated expectations,

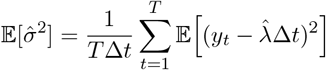

and

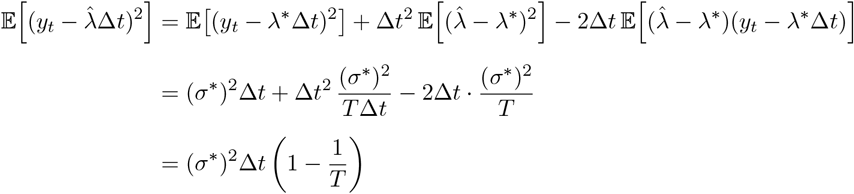

Thus,

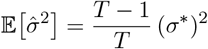

This is known as Bessel effect and can explain a noticeable downward bias when each replay event contains only a handful of time bins.

#### Extension to switching HMM

We consider a switching HMM with a discrete regime *c*_*t*_ that controls the transition kernel, and only one regime (noted as *s* = ⋆) has drift-diffusion parameters (*λ, σ*) while the others are fixed (i.e., stationary or uniform). Extending the above bias analysis should be relatively straightforward.

In the above HMM setup, the score for *λ* and *σ* come entirely from the transition terms and use pairwise marginals,

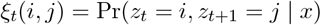

In the switching HMM, the log-joint includes the regime chain and the transition kernel is 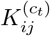. The score for *θ* = (*λ, σ*) only accrues when *c*_*t*_ = ⋆. Practically, we can replace *ξ*_*t*_(*i, j*) by regime-conditioned pairwise marginals,

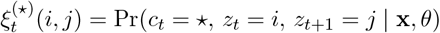

and ignore transitions from other regimes in the derivative, because their kernels are not relevant to (*λ, σ*).

A subsequent change in derivation is that HMM formulas become weighted by the posterior probability that the ⋆ regime is active at time *t*:

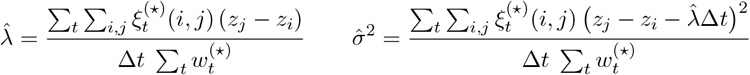

with the regime-posterior weight,

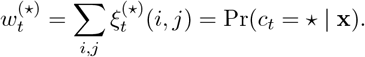

Similar to HMM case, for 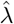. under the correctly specified switching model and evaluate the posteriors at the true parameters (*λ*^∗^, (*σ*^∗^)^2^),

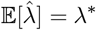

For 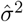, the total time step *T* in above HMM expressions should be replaced by Kish’s effective sample size,

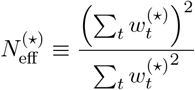

The expectation of (*σ*^∗^)^2^ estimator becomes:

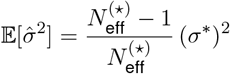

I.e., 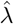 is unbiased and the finite-sample downward bias for 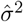, with *T* replaced by 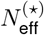.

## References

[1] Brad E Pfeiffer and David J Foster. Hippocampal place-cell sequences depict future paths to remembered goals. Nature, 497(7447):74–79, 2013.

[2] Kamran Diba and György Buzsáki. Forward and reverse hippocampal place-cell sequences during ripples. Nature neuroscience, 10(10):1241–1242, 2007.

[3] David J Foster and Matthew A Wilson. Reverse replay of behavioural sequences in hippocampal place cells during the awake state. Nature, 440(7084):680–683, 2006.

[4] R Ellen Ambrose, Brad E Pfeiffer, and David J Foster. Reverse replay of hippocampal place cells is uniquely modulated by changing reward. Neuron, 91(5):1124–1136, 2016.

[5] Matthew A Wilson and Bruce L McNaughton. Reactivation of hippocampal ensemble memories during sleep. Science, 265(5172):676–679, 1994.

[6] William E Skaggs and Bruce L McNaughton. Replay of neuronal firing sequences in rat hippocampus during sleep following spatial experience. Science, 271(5257):1870–1873, 1996.

[7] Zoltán Nádasdy, Hajime Hirase, András Czurkó, Jozsef Csicsvari, and György Buzsáki. Replay and time compression of recurring spike sequences in the hippocampus. Journal of neuroscience, 19(21):9497–9507, 1999.

[8] Albert K Lee and Matthew A Wilson. Memory of sequential experience in the hippocampus during slow wave sleep. Neuron, 36(6):1183–1194, 2002.

[9] Mattias P Karlsson and Loren M Frank. Awake replay of remote experiences in the hippocampus. Nature neuroscience, 12(7):913–918, 2009.

[10] Adrien Peyrache, Mehdi Khamassi, Karim Benchenane, Sidney I Wiener, and Francesco P Battaglia. Replay of rule-learning related neural patterns in the prefrontal cortex during sleep. Nature neuroscience, 12(7):919–926, 2009.

[11] Justin D Shin, Wenbo Tang, and Shantanu P Jadhav. Dynamics of awake hippocampal-prefrontal replay for spatial learning and memory-guided decision making. Neuron, 104(6):1110–1125, 2019.

[12] Karola Kaefer, Michele Nardin, Karel Blahna, and Jozsef Csicsvari. Replay of behavioral sequences in the medial prefrontal cortex during rule switching. Neuron, 106(1):154–165, 2020.

[13] Daoyun Ji and Matthew A Wilson. Coordinated memory replay in the visual cortex and hippocampus during sleep. Nature neuroscience, 10(1):100–107, 2007.

[14] Yifan Yang, David A Leopold, Jeff H Duyn, and Xiao Liu. Hippocampal replay sequence governed by spontaneous brain-wide dynamics. PNAS nexus, 3(4):pgae078, 2024.

[15] Anna C Schapiro, Elizabeth A McDevitt, Timothy T Rogers, Sara C Mednick, and Kenneth A Norman. Human hippocampal replay during rest prioritizes weakly learned information and predicts memory performance. Nature communications, 9(1):3920, 2018.

[16] Yitzhak Norman, Erin M Yeagle, Simon Khuvis, Michal Harel, Ashesh D Mehta, and Rafael Malach. Hippocampal sharp-wave ripples linked to visual episodic recollection in humans. Science, 365(6454):eaax1030, 2019.

[17] Yunzhe Liu, Raymond J Dolan, Zeb Kurth-Nelson, and Timothy EJ Behrens. Human replay spontaneously reorganizes experience. Cell, 178(3):640–652, 2019.

[18] Yunzhe Liu, Marcelo G Mattar, Timothy EJ Behrens, Nathaniel D Daw, and Raymond J Dolan. Experience replay is associated with efficient nonlocal learning. Science, 372(6544):eabf1357, 2021.

[19] Yunzhe Liu, Matthew M Nour, Nicolas W Schuck, Timothy EJ Behrens, and Raymond J Dolan. Decoding cognition from spontaneous neural activity. Nature Reviews Neuroscience, 23(4):204–214, 2022.

[20] David J Foster. Replay comes of age. Annual review of neuroscience, 40(1):581–602, 2017.

[21] Laura Lee Colgin. Five decades of hippocampal place cells and eeg rhythms in behaving rats. Journal of Neuroscience, 40(1):54–60, 2020.

[22] Kimberly L Stachenfeld, Matthew M Botvinick, and Samuel J Gershman. The hippocampus as a predictive map. Nature neuroscience, 20(11):1643–1653, 2017.

[23] Zhe Sage Chen and Matthew A Wilson. How our understanding of memory replay evolves. Journal of Neurophysiology, 129(3):552–580, 2023.

[24] David Tingley and Adrien Peyrache. On the methods for reactivation and replay analysis. Philosophical Transactions of the Royal Society B, 375(1799):20190231, 2020.

[25] Eric L Denovellis, Anna K Gillespie, Michael E Coulter, Marielena Sosa, Jason E Chung, Uri T Eden, and Loren M Frank. Hippocampal replay of experience at real-world speeds. Elife, 10:e64505, 2021.

[26] Emma L Krause and Jan Drugowitsch. A large majority of awake hippocampal sharp-wave ripples feature spatial trajectories with momentum. Neuron, 110(4):722–733, 2022.

[27] Daniel C McNamee, Kimberly L Stachenfeld, Matthew M Botvinick, and Samuel J Gershman. Flexible modulation of sequence generation in the entorhinal–hippocampal system. Nature neuroscience, 24(6):851–862, 2021.

[28] Riccardo Barbieri, Matthew A Wilson, Loren M Frank, and Emery N Brown. An analysis of hippocampal spatio-temporal representations using a bayesian algorithm for neural spike train decoding. IEEE transactions on neural systems and rehabilitation engineering, 13(2):131–136, 2005.

[29] Marc Box, Matt W Jones, and Nick Whiteley. A hidden markov model for decoding and the analysis of replay in spike trains. Journal of computational neuroscience, 41(3):339–366, 2016.

[30] Kourosh Maboudi, Etienne Ackermann, Laurel Watkins de Jong, Brad E Pfeiffer, David Foster, Kamran Diba, and Caleb Kemere. Uncovering temporal structure in hippocampal output patterns. Elife, 7:e34467, 2018.

[31] Federico Stella, Peter Baracskay, Joseph O’Neill, and Jozsef Csicsvari. Hippocampal reactivation of random trajectories resembling brownian diffusion. Neuron, 102(2):450–461, 2019.

[32] George Dragoi and Susumu Tonegawa. Preplay of future place cell sequences by hippocampal cellular assemblies. Nature, 469(7330):397–401, 2011.

[33] Brad E Pfeiffer and David J Foster. Autoassociative dynamics in the generation of sequences of hippocampal place cells. Science, 349(6244):180–183, 2015.

[34] Kechen Zhang, Iris Ginzburg, Bruce L McNaughton, and Terrence J Sejnowski. Interpreting neuronal population activity by reconstruction: unified framework with application to hippocampal place cells. Journal of neurophysiology, 79(2):1017–1044, 1998.

[35] Xiaojing Wu and David J Foster. Hippocampal replay captures the unique topological structure of a novel environment. Journal of Neuroscience, 34(19):6459–6469, 2014.

[36] Leonard E Baum, Ted Petrie, George Soules, and Norman Weiss. A maximization technique occurring in the statistical analysis of probabilistic functions of markov chains. The annals of mathematical statistics, 41(1):164–171, 1970.

[37] Lawrence R Rabiner. A tutorial on hidden markov models and selected applications in speech recognition. Proceedings of the IEEE, 77(2):257–286, 2002.

[38] Zhe Chen, Fabian Kloosterman, Emery N Brown, and Matthew A Wilson. Uncovering spatial topology represented by rat hippocampal population neuronal codes. Journal of computational neuroscience, 33(2):227–255, 2012.

[39] Zhe Chen. An overview of bayesian methods for neural spike train analysis. Computational intelligence and neuroscience, 2013(1):251905, 2013.

[40] Scott W Linderman, Matthew J Johnson, Matthew A Wilson, and Zhe Chen. A bayesian nonparametric approach for uncovering rat hippocampal population codes during spatial navigation. Journal of neuroscience methods, 263:36–47, 2016.

[41] Scott Linderman, Matthew Johnson, Andrew Miller, Ryan Adams, David Blei, and Liam Paninski. Bayesian learning and inference in recurrent switching linear dynamical systems. In Artificial intelligence and statistics, pages 914–922. PMLR, 2017.

[42] Joshua Glaser, Matthew Whiteway, John P Cunningham, Liam Paninski, and Scott Linderman. Recurrent switching dynamical systems models for multiple interacting neural populations. Advances in neural information processing systems, 33:14867–14878, 2020.

[43] Felix Pei, Joel Ye, David Zoltowski, Anqi Wu, Raeed H Chowdhury, Hansem Sohn, Joseph E O’Doherty, Krishna V Shenoy, Matthew T Kaufman, Mark Churchland, et al. Neural latents benchmark’21: evaluating latent variable models of neural population activity. arXiv preprint arXiv:2109.04463, 2021.

[44] Carles Balsells-Rodas, Yixin Wang, and Yingzhen Li. On the identifiability of switching dynamical systems. arXiv preprint arXiv:2305.15925, 2023.

[45] Yonghong An, Yingyao Hu, Johns Hopkins, and Matt Shum. Identifiability and inference of hidden markov models. Technical report, Technical report, 2013.

[46] Delia Silva, Ting Feng, and David J Foster. Trajectory events across hippocampal place cells require previous experience. Nature neuroscience, 18(12):1772–1779, 2015.

[47] Masahiro Takigawa, Marta Huelin Gorriz, Margot Tirole, and Daniel Bendor. Evaluating hippocampal replay without a ground truth. ELife, 13:e85635, 2024.

[48] Alexander Lex, Nils Gehlenborg, Hendrik Strobelt, Romain Vuillemot, and Hanspeter Pfister. Upset: visualization of intersecting sets. IEEE transactions on visualization and computer graphics, 20(12):1983–1992, 2014.

[49] Thomas J Davidson, Fabian Kloosterman, and Matthew A Wilson. Hippocampal replay of extended experience. Neuron, 63(4):497–507, 2009.

[50] Usman Farooq, Jeremie Sibille, Kefei Liu, and George Dragoi. Strengthened temporal coordination within pre-existing sequential cell assemblies supports trajectory replay. Neuron, 103(4):719–733, 2019.

[51] Michael N Shadlen and William T Newsome. Neural basis of a perceptual decision in the parietal cortex (area lip) of the rhesus monkey. Journal of neurophysiology, 86(4):1916–1936, 2001.

[52] Jamie D Roitman and Michael N Shadlen. Response of neurons in the lateral intraparietal area during a combined visual discrimination reaction time task. Journal of neuroscience, 22(21):9475–9489, 2002.

[53] Joshua I Gold and Michael N Shadlen. The neural basis of decision making. Annu. Rev. Neurosci., 30(1):535–574, 2007.

[54] Wenbo Tang, Justin D Shin, Loren M Frank, and Shantanu P Jadhav. Hippocampal-prefrontal reactivation during learning is stronger in awake compared with sleep states. Journal of Neuroscience, 37(49):11789–11805, 2017.

[55] Justin D Shin, Wenbo Tang, and Shantanu P Jadhav. Dynamics of awake hippocampal-prefrontal replay for spatial learning and memory-guided decision making. Neuron, 104(6):1110–1125, 2019.

[56] Jordan Breffle, Hannah Germaine, Justin D Shin, Shantanu P Jadhav, and Paul Miller. Intrinsic dynamics of randomly clustered networks generate place fields and preplay of novel environments. Elife, 13:RP93981, 2024.

[57] Andres D Grosmark and György Buzsáki. Diversity in neural firing dynamics supports both rigid and learned hippocampal sequences. Science, 351(6280):1440–1443, 2016.

[58] George Dragoi and György Buzsáki. Temporal encoding of place sequences by hippocampal cell assemblies. Neuron, 50(1):145–157, 2006.

[59] David J Foster and Matthew A Wilson. Hippocampal theta sequences. Hippocampus, 17(11):1093–1099, 2007.

[60] Chenguang Zheng, Ernie Hwaun, Carlos A Loza, and Laura Lee Colgin. Hippocampal place cell sequences differ during correct and error trials in a spatial memory task. Nature communications, 12(1):3373, 2021.

[61] Andrew M Wikenheiser and A David Redish. Hippocampal theta sequences reflect current goals. Nature neuroscience, 18(2):289–294, 2015.

[62] Kenneth Kay, Jason E Chung, Marielena Sosa, Jonathan S Schor, Mattias P Karlsson, Margaret C Larkin, Daniel F Liu, and Loren M Frank. Constant sub-second cycling between representations of possible futures in the hippocampus. Cell, 180(3):552–567, 2020.

[63] Abraham Z Vollan, Richard J Gardner, May-Britt Moser, and Edvard I Moser. Left–right-alternating theta sweeps in entorhinal–hippocampal maps of space. Nature, 639(8056):995–1005, 2025.

[64] Long-Ji Lin. Self-improving reactive agents based on reinforcement learning, planning and teaching. Machine learning, 8(3):293–321, 1992.

[65] Volodymyr Mnih, Koray Kavukcuoglu, David Silver, Alex Graves, Ioannis Antonoglou, Daan Wierstra, and Martin Riedmiller. Playing atari with deep reinforcement learning. arXiv preprint arXiv:1312.5602, 2013.

[66] Volodymyr Mnih, Koray Kavukcuoglu, David Silver, Andrei A Rusu, Joel Veness, Marc G Bellemare, Alex Graves, Martin Riedmiller, Andreas K Fidjeland, Georg Ostrovski, et al. Human-level control through deep reinforcement learning. nature, 518(7540):529–533, 2015.

[67] German I Parisi, Ronald Kemker, Jose L Part, Christopher Kanan, and Stefan Wermter. Continual lifelong learning with neural networks: A review. Neural networks, 113:54–71, 2019.

[68] Matthias De Lange, Rahaf Aljundi, Marc Masana, Sarah Parisot, Xu Jia, Aleš Leonardis, Gregory Slabaugh, and Tinne Tuytelaars. A continual learning survey: Defying forgetting in classification tasks. IEEE transactions on pattern analysis and machine intelligence, 44(7):3366–3385, 2021.

[69] Gido M Van de Ven, Tinne Tuytelaars, and Andreas S Tolias. Three types of incremental learning. Nature Machine Intelligence, 4(12):1185–1197, 2022.

[70] Tom Schaul, John Quan, Ioannis Antonoglou, and David Silver. Prioritized experience replay. arXiv preprint arXiv:1511.05952, 2015.

[71] David Rolnick, Arun Ahuja, Jonathan Schwarz, Timothy Lillicrap, and Gregory Wayne. Experience replay for continual learning. Advances in neural information processing systems, 32, 2019.

[72] Hanul Shin, Jung Kwon Lee, Jaehong Kim, and Jiwon Kim. Continual learning with deep generative replay. Advances in neural information processing systems, 30, 2017.

[73] Gido M Van de Ven, Hava T Siegelmann, and Andreas S Tolias. Brain-inspired replay for continual learning with artificial neural networks. Nature communications, 11(1):4069, 2020.

[74] Rui Gao and Weiwei Liu. Ddgr: Continual learning with deep diffusion-based generative replay. In International Conference on Machine Learning, pages 10744–10763. PMLR, 2023.

[75] Raia Hadsell, Dushyant Rao, Andrei A Rusu, and Razvan Pascanu. Embracing change: Continual learning in deep neural networks. Trends in cognitive sciences, 24(12):1028–1040, 2020.

[76] Lennart Wittkuhn, Samson Chien, Sam Hall-McMaster, and Nicolas W Schuck. Replay in minds and machines. Neuroscience & Biobehavioral Reviews, 129:367–388, 2021.

[77] Emily A Jones, Anna K Gillespie, Seo Yeon Yoon, Loren M Frank, and Yadong Huang. Early hippocampal sharp-wave ripple deficits predict later learning and memory impairments in an alzheimer’s disease mouse model. Cell reports, 29(8):2123–2133, 2019.

[78] Valérie Ego-Stengel and Matthew A Wilson. Disruption of ripple-associated hippocampal activity during rest impairs spatial learning in the rat. Hippocampus, 20(1):1–10, 2010.

[79] Antonio Fernández-Ruiz, Azahara Oliva, Eliezyer Fermino de Oliveira, Florbela Rocha-Almeida, David Tingley, and György Buzsáki. Long-duration hippocampal sharp wave ripples improve memory. Science, 364(6445):1082–1086, 2019.

[80] Sarah Shipley, Marco P Abrate, Robin Hayman, Dennis Chan, and Caswell Barry. Disordered hippocampal reactivations predict spatial memory deficits in a mouse model of alzheimer’s disease. bioRxiv, pages 2024–11, 2024.

[81] Igor Gridchyn, Philipp Schoenenberger, Joseph O’Neill, and Jozsef Csicsvari. Assembly-specific disruption of hippocampal replay leads to selective memory deficit. Neuron, 106(2):291–300, 2020.

[82] Michael E Coulter, Anna K Gillespie, Joshua Chu, Eric L Denovellis, Trevor Thai K Nguyen, Daniel F Liu, Katherine Wadhwani, Baibhav Sharma, Kevin Wang, Xinyi Deng, et al. Closed-loop modulation of remote hippocampal representations with neurofeedback. Neuron, 113(6):949–961, 2025.

## References

[1] Kechen Zhang, Iris Ginzburg, Bruce L McNaughton, and Terrence J Sejnowski. Interpreting neuronal population activity by reconstruction: unified framework with application to hippocampal place cells. Journal of neurophysiology, 79(2):1017–1044, 1998.

[2] Delia Silva, Ting Feng, and David J Foster. Trajectory events across hippocampal place cells require previous experience. Nature neuroscience, 18(12):1772–1779, 2015.

[3] Masahiro Takigawa, Marta Huelin Gorriz, Margot Tirole, and Daniel Bendor. Evaluating hippocampal replay without a ground truth. ELife, 13:e85635, 2024.

[4] Usman Farooq, Jeremie Sibille, Kefei Liu, and George Dragoi. Strengthened temporal coordination within pre-existing sequential cell assemblies supports trajectory replay. Neuron, 103(4):719–733, 2019.

[5] Matthijs AA van der Meer, Caleb Kemere, and Kamran Diba. Progress and issues in second-order analysis of hippocampal replay. Philosophical Transactions of the Royal Society B, 375(1799):20190238, 2020.

[6] Brad E Pfeiffer and David J Foster. Autoassociative dynamics in the generation of sequences of hippocampal place cells. Science, 349(6244):180–183, 2015.

